# Active repression of muscle fate preserves neural lineage identity during cerebellum development

**DOI:** 10.64898/2026.05.13.725001

**Authors:** Nurunnabi Mirja Shaikh, Venkata Thulabandu, Akira Inoue, Joshua Paré, Jackie Norrie, Qiong Zhang, Beisi Xu, Xinwei Cao

## Abstract

Cell fate commitment is commonly thought to entail progressive restriction of developmental potential, enforced by passive, heterochromatin-based silencing of alternative lineage programs. Here we show that maintenance of neural identity during cerebellum development instead requires active repression of a starkly divergent fate by the TEAD–INSM1 transcriptional complex. Loss of TEAD1/2 or INSM1 activates the myogenic master regulator *Myod1*, resulting in neural cells acquiring transcriptional, structural, and metabolic features of skeletal muscle cells. Deletion of *Myod1* fully suppresses neural-to-muscle conversion while partially rescuing neural developmental defects. Our results uncover a latent alternative lineage during neurodevelopment and a surprising role for sequence-specific transcription factors in enforcing lineage boundaries, including those previously thought essentially unbreachable, with implications for understanding aberrant differentiation in disease contexts and cell-type evolution.

---

Metazoan development involves a series of hierarchical, unidirectional cell fate choices, beginning with germ layer separation and followed by acquisition of increasingly refined cell identities. When one fate is chosen, alternative fates are repressed or silenced to ensure unambiguous and stable cell identity. Although it is now well recognized that cells can change lineages (*1–4*), particularly upon forced expression of lineage–instructive transcription factors (TFs), a general principle is that the greater the developmental distance, the harder it is to convert (*2*). Accordingly, transdifferentiation in vivo does not cross germ layers (*3–5*) except for rare disease contexts, underscoring exceptionally strong barriers between lineages arising from different germ layers. The central role of cross-antagonistic interactions between lineage–instructive TFs during cell fate commitment is well established (*2*), as is the importance of heterochromatin states and three-dimensional genome organization in silencing inappropriate lineage programs (*6–10*). But by contrast, the role of sequence-specific TFs in actively repressing developmentally distant lineages in vivo is almost completely unexplored.

The TEAD family of TFs (TEAD1–4 in mammals) are the DNA-binding factors of the Hippo pathway, where they recruit the transcriptional coactivators YAP and TAZ to regulate cell proliferation and tissue growth (*11*) and, in select developmental contexts, binary cell-fate decisions (*12, 13*). Consistent with the fact that TEAD predates YAP in evolution (*14, 15*), it can function outside canonical YAP/TAZ-mediated Hippo signaling through interactions with alternative cofactors. In particular, TEAD interacts with INSM1 (insulinoma-associated 1) (*16, 17*), a SNAG (Snail/Gfi-1)-domain containing zinc-finger TF (*18*), to regulate the differentiation of specific neurons in *C. elegans* (*19*), maintain the differentiated state of medulla neurons in *Drosophila* optic lobe (*16*), and promote developmental progression of neural progenitors in the mammalian ventral telencephalon (*20*). As TEAD factors are expressed more broadly than YAP/TAZ across the developing nervous system (*21, 22*), we reasoned that TEAD may exert Hippo-independent roles more widely during neural development. We focused on the cerebellum—where YAP and TAZ play only minor roles (*23*)—and uncovered an unexpected and remarkable requirement for TEAD and INSM1 in maintaining neural lineage identity by actively repressing a developmentally distant alternative fate.

## TEAD1/2 and INSM1 are coexpressed in cerebellar granule neuron precursors

The most abundant cells in the cerebellum—and indeed in the entire brain—cerebellar granule neurons, are produced by granule neuron precursors (GNPs). GNPs originate in the upper rhombic lip (uRL) of the hindbrain and form a secondary germinal zone, the external granule layer (EGL), during embryonic development (*24*) (Fig. 1A). In mice, GNPs in the EGL proliferate extensively during the first two postnatal weeks, producing large numbers of granule neurons and driving rapid cerebellar growth. Postmitotic granule neurons then migrate inward and settle in the internal granule layer (IGL), where they mature.

**Fig. 1.**
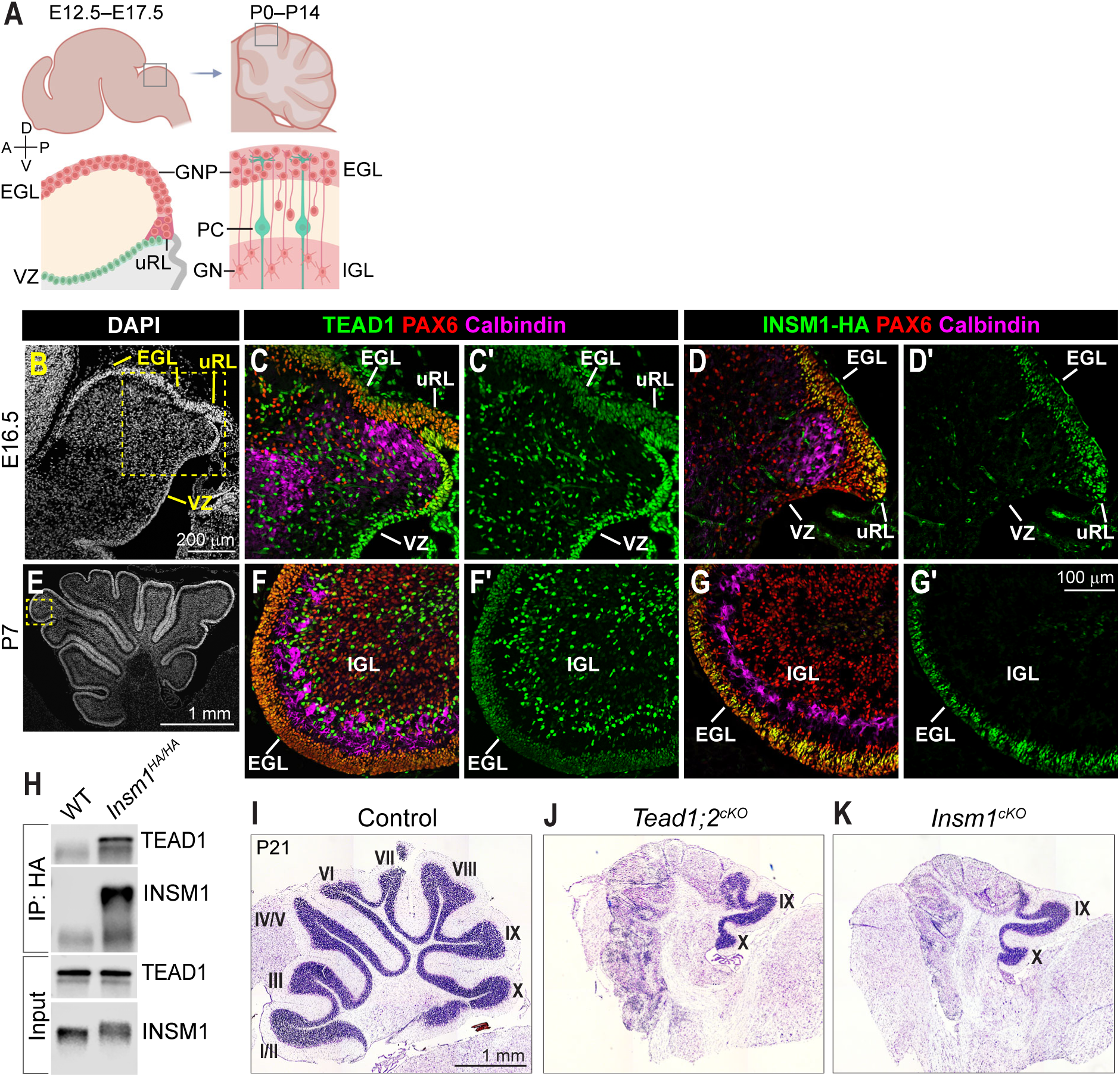
TEAD1/2 and INSM1 are essential for cerebellum development. (**A**) A diagram of mouse cerebellum development, depicting granule neuron precursor (GNPs) arising from the upper rhombic lip (uRL) and forming the external granule layer (EGL) during embryonic day (E) 12.5–17.5, proliferating in the EGL during postnatal day (P) 0–14, before migrating to the internal granule layer (IGL) and differentiating into granule neurons (GN). (**B**) DAPI nuclear staining of an E16.5 wild-type (WT) mouse cerebellum midsagittal section. (**C** and **C′**) Co-immunostaining of E16.5 WT cerebellum for TEAD1, the granule neuron lineage marker PAX6, and the Purkinje cell (PC) marker Calbindin. (**D** and **D′**) Co-immunostaining of E16.5 *Insm1^HA/HA^* cerebellum for HA (INSM1), PAX6, and Calbindin. (**E**) DAPI nuclear staining of a P7 WT cerebellum midsagittal section. (**F** and **F′**) Co-immunostaining of P7 WT cerebellum for TEAD1, PAX6, and Calbindin. (**G** and **G′**) Co-immunostaining of P7 *Insm1^HA/HA^* cerebellum for HA (INSM1), PAX6, and Calbindin. (**H**) Co-immunoprecipitation (co-IP) of TEAD1 by an anti-HA antibody from P7 *Insm1^HA/HA^*, but not WT (negative control), cerebellar nuclear extracts. (**I** to **K**) Nissl staining of P21 control and mutant cerebella near-vermis sections with lobule numbers labeled. VZ, ventricular zone; Boxed regions in B and E are shown at higher magnification in C and F, respectively. Illustration in A was created partly using BioRender.

To define the role of TEAD factors during cerebellum development, we first examined their expression patterns. Co-immunostaining of developing cerebellum with antibodies against TEAD1, PAX6—a marker of the granule neuron lineage, and Calbindin—a marker of Purkinje cells, revealed that TEAD1 was expressed in ventricular zone (VZ) neural progenitors, GNPs in the uRL and EGL, and BLBP (FABP7; fatty acid binding protein 7)-labeled glial cells in the IGL, but not in Purkinje cells or granule neurons in the IGL (Fig. 1, B to C′, E to F′; fig. S1, A and B). *Tead2* mRNA was also detected in the VZ, uRL, and EGL at embryonic day 15.5 (E15.5; fig. S1C) (*25*).

Given prior evidence that TEAD acts with INSM1 in other neurodevelopmental contexts (*16, 19, 20, 26*), we next examined whether these factors are coexpressed in the developing cerebellum. A previous study reported *Insm1* mRNA expression in the EGL but not in the IGL (*27*). To further define INSM1 expression, we generated an *Insm1^HA^* knock-in mouse line by fusing a hemagglutinin (HA) tag to the C-terminus of endogenous INSM1. *Insm1^HA/HA^* mice are viable and fertile, whereas *Insm1^−/−^* mice die before birth (*28, 29*), indicating that the HA-tag does not disrupt INSM1 function. The cerebellum of *Insm1^HA/HA^* mice was morphologically indistinguishable from that of wild-type (WT) mice (fig. S1, D and E). Co-immunostaining with antibodies against HA (for INSM1), PAX6, and Calbindin showed that INSM1 was expressed in GNPs in the uRL and EGL, but not in VZ progenitors, Purkinje cells, or granule neurons in the IGL (Fig. 1, D, D′, G, and G′). Thus, TEAD1/2 and INSM1 are coexpressed in GNPs during cerebellum development. We further confirmed the interaction between TEAD1 and INSM1 in cerebellar nuclear extracts by co-immunoprecipitation (Fig. 1H; fig. S1F).

## TEAD1/2 and INSM1 are essential for cerebellum development

To determine the roles of TEAD and INSM1 in GNPs, we generated conditional knockout (cKO) mice using an *Atoh1-Cre* line (*30*), which targets GNPs in anterior and central lobules but with only sparse activity in lobules IX and X (*31*) (fig. S1, G to I′). In *Tead1^F/F^;Tead2^F/F^;Atoh1-Cre* (*Tead1;2^cKO^*) mice, TEAD1 was undetectable in the EGL of anterior and central lobules at postnatal day 0 (P0; fig. S1, G to G′′), confirming efficient deletion in GNPs. *Tead1^cKO^* cerebella exhibited reduced IGL thickness in lobules I–VII (fig. S1J), consistent with a previous report (*32*). *Tead2^cKO^* and *Insm1^F/+^;Atoh1-Cre* (*Insm1^Het^*) cerebella displayed no overt morphological abnormalities (fig. S1, K and L). Strikingly, both *Tead1;2^cKO^* and *Insm1^cKO^* cerebella showed severe hypoplasia and malformation (Fig. 1, I to K).

The morphological defects of mutant cerebella were already evident at P3 (Fig. 2, A to C). Although a nuclear-dense EGL remained recognizable in mutant cerebella, TAG1 (Contactin 2), which labels newly differentiated granule neurons in the inner EGL, and NeuN (Rbfox3), which marks more mature granule neurons in the IGL, were significantly reduced in regions corresponding to anterior and central lobules in both mutant lines compared with no-*Cre* controls (Fig. 2, A to C′, J, and K). At P7, cerebellar morphology deteriorated further: the external, EGL-like layer was occupied by NeuN^+^ cells, and both TAG1^+^ and NeuN^+^ cells remained significantly reduced (Fig. 2, D to F′, J, and K). In addition, significantly higher fractions of GNPs exited the cell cycle between P2 and P3 in both mutant lines (Fig. 2, G to I, and L). Cell death was also increased in both mutants (fig. S2), likely contributing to cerebellar malformation. Together, these results indicate that TEAD1/2 and INSM1 are essential for the proliferation, differentiation, and survival of cerebellar GNPs.

**Fig. 2.**
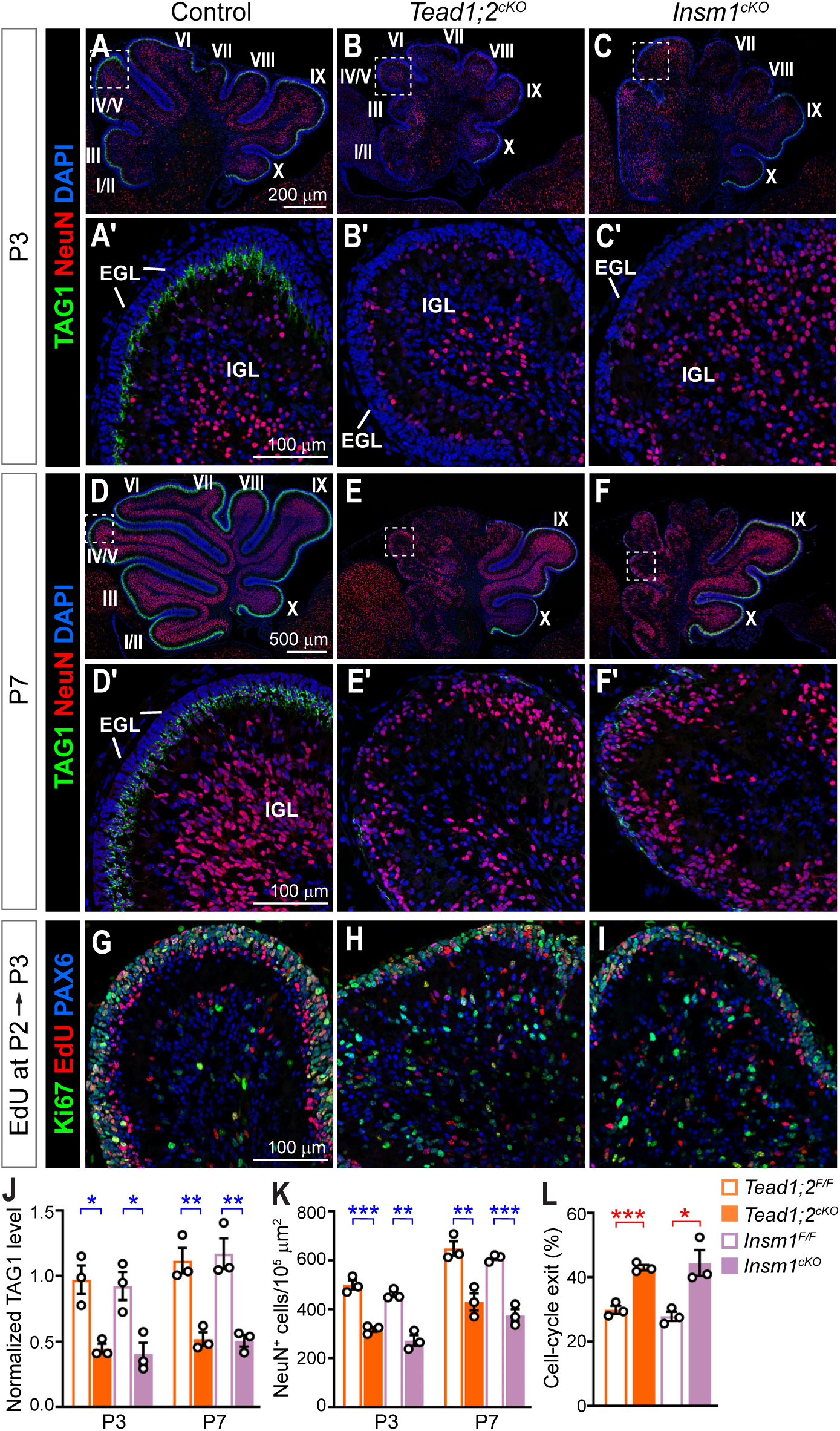
TEAD1/2 and INSM1 are essential for GNP differentiation and proliferation. (**A** to **F′**) Co-immunostaining of P3 and P7 cerebella for the newborn neuron marker TAG1 and the more mature neuron marker NeuN. (**G** to **I**) Co-immunostaining of P3 cerebella that were labeled with 5-ethynyl-2′-deoxyuridine (EdU) at P2. (**J**) Quantification of TAG1 immunofluorescence signals per unit area in lobules III–V normalized to the signals in lobule IX in the same section. (**K**) Quantifications of the number of NeuN^+^ cells in lobules III–V. (**L**) GNP cell cycle exit index calculated as EdU^+^PAX6^+^Ki67^−^ cells/EdU^+^PAX6^+^ cells. Each data point represents an individual animal. Values are mean ± SEM. Unpaired two-tailed *t*-test; *, *P* < 0.05; **, *P* < 0.01; ***, *P* < 0.001. EGL, external granule layer; IGL, internal granule layer. Boxed regions in A to F are shown at higher magnification in A′ to F′.

## Loss of TEAD1/2 or INSM1 downregulates neural genes and upregulates muscle genes

To capture the early and potentially direct effects of TEAD1/2 and INSM1 loss on GNP gene expression, we performed RNA sequencing (RNA-seq) on P2 cerebella, a stage at which mutant phenotypes were relatively subtle. Differentially expressed genes identified in *Tead1;2^cKO^* and *Insm1^cKO^* cerebella (each compared with littermate no-*Cre* controls) overlapped significantly (Fig. 3A). Consistent with our immunostaining results, genes involved in neural development were downregulated. Unexpectedly, genes associated with skeletal muscle development—including myogenic regulatory factors (MRFs) *Myod1* and *Myogenin* (*Myog*), and muscle structural genes such as *Actc1* and *Ttn*—were upregulated in both mutant lines (Fig. 3B; fig. S3A; table S1).

**Fig. 3.**
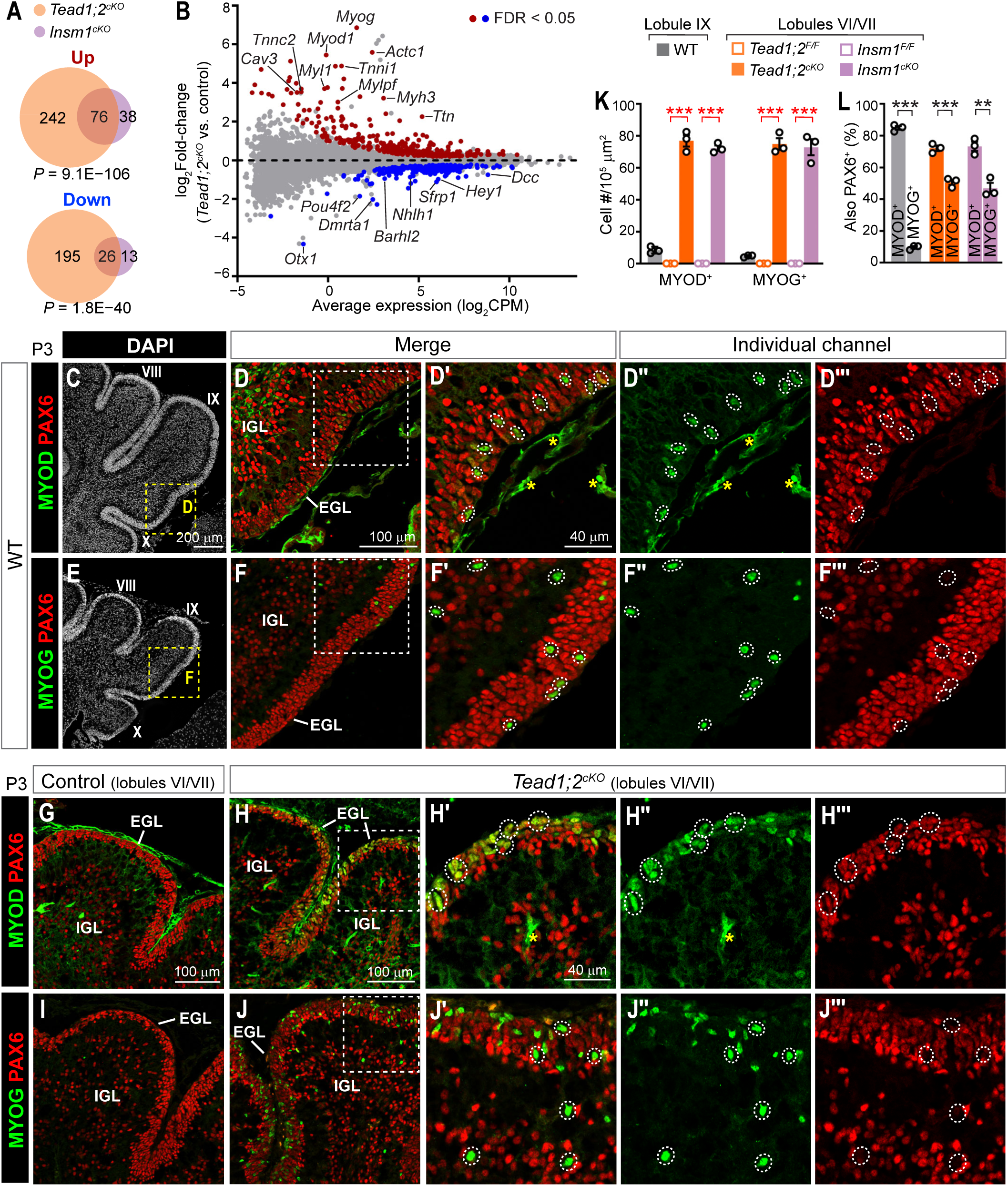
Loss of TEAD1/2 or INSM1 upregulates muscle genes. (**A**) Venn diagram showing the numbers and overlap of significantly up- and down-regulated genes (false discovery rate (FDR) < 0.05, no fold-change cutoff) between *Tead1;2^cKO^* and *Insm1^cKO^* cerebella, each compared with corresponding littermate controls, based on RNA-sequencing. Overlap *P* value, hypergeometric test. (**B**) MA plot showing gene expression changes in *Tead1;2^cKO^* cerebella. (**C** to **F′′′**) Co-immunostaining of P3 WT cerebellum for PAX6 and MYOD or myogenin (MYOG). Asterisks, nonspecific staining. (**G** to **J′′′**) Co-immunostaining of P3 control and *Tead1;2^cKO^* cerebella for PAX6 and MYOD or MYOG. (**K**) Quantifications of the number of MYOD^+^ or MYOG^+^ cells in lobule IX of WT cerebella or lobules VI/VII of control and mutant cerebella. (**L**) Quantifications of the fraction of MYOD^+^ or MYOG^+^ cells that were also PAX6^+^. Each data point represents an individual animal. Values are mean ± SEM. Unpaired two-tailed *t*-test; **, *P* < 0.01; ***, *P* < 0.001. Boxed regions in C and E are shown at higher magnification in D and F, respectively. Boxed regions in D, F, H, and J are enlarged in D′, F′, H′, and J′. Dotted circles highlight MYOD^+^ or MYOG^+^ cells. EGL, external granule layer; IGL, internal granule layer.

MRFs, which include MYOD, Myogenic factor 5 (MYF5), MYF6, and MYOG, are basic helix-loop-helix (bHLH) TFs that orchestrate skeletal muscle development. MYOD, MYF5, and MYF6 direct stem cells to the skeletal muscle lineage, and MYOG—along with MYOD and MYF6—activates the myogenic differentiation program (*33*). To validate the RNA-seq results at the protein level, we performed immunostaining for MYOD and MYOG. Consistent with a previous report of sporadic MYOD^+^ cells in the EGL of WT cerebellum (*34*), we also detected MYOD^+^ cells in P3 cerebella of both no-*Cre* control mice and WT mice without genetically modified alleles. These cells were largely restricted to lobule IX (Fig. 3, C to D′′′, and K; fig. S3, B to B′′′). They were predominantly located in the EGL and most were positive for PAX6, indicating that they were GNPs; however, their PAX6 levels were generally lower than those of neighboring MYOD^−^ cells (Fig. 3, D′ to D′′′, and L). We also detected MYOG^+^ cells in WT and control cerebella; they were largely restricted to lobule IX and were often negative or weakly positive for PAX6 (Fig. 3, E to F′′′, K, and L).

In both *Tead1;2^cKO^* and *Insm1^cKO^* cerebella, MYOD^+^ and MYOG^+^ cells were markedly increased at P3, particularly along the surface of lobules VI–VIII (Fig. 3, G to K; fig. S3, D to D′′′). In mutant as well as WT cerebella, the fraction of MYOG^+^ cells that were also positive for PAX6 was significantly lower than that of MYOD^+^ cells (Fig. 3L), suggesting a progressive loss of granule neuron lineage identity.

## Loss of TEAD1/2 or INSM1 induces complete neural-to-muscle lineage conversion

Next, we examined muscle structural proteins, including troponin C2 (TNNC2), myosin heavy chain (MYH), and myosin light chain 1, etc. Remarkably, muscle protein–positive cells were abundant in the P7 cerebellum of both *Tead1;2^cKO^* and *Insm1^cKO^* mice and persisted at P21 (Fig. 4, A to B′, E, and E′; fig. S4), with some even expressing the adult fast muscle myosin MYH1 at P21 (fig. S4, Q and R) (*35*). In contrast, despite the presence of MYOD^+^ and MYOG^+^ cells in control cerebella, muscle protein–positive cells were exceedingly rare (fig. S4, A, D, G, and J), suggesting that TEAD1/2 and INSM1 not only suppress *Myod1* and *Myog* expression but also restrain muscle differentiation downstream of MYOD/MYOG.

**Fig. 4.**
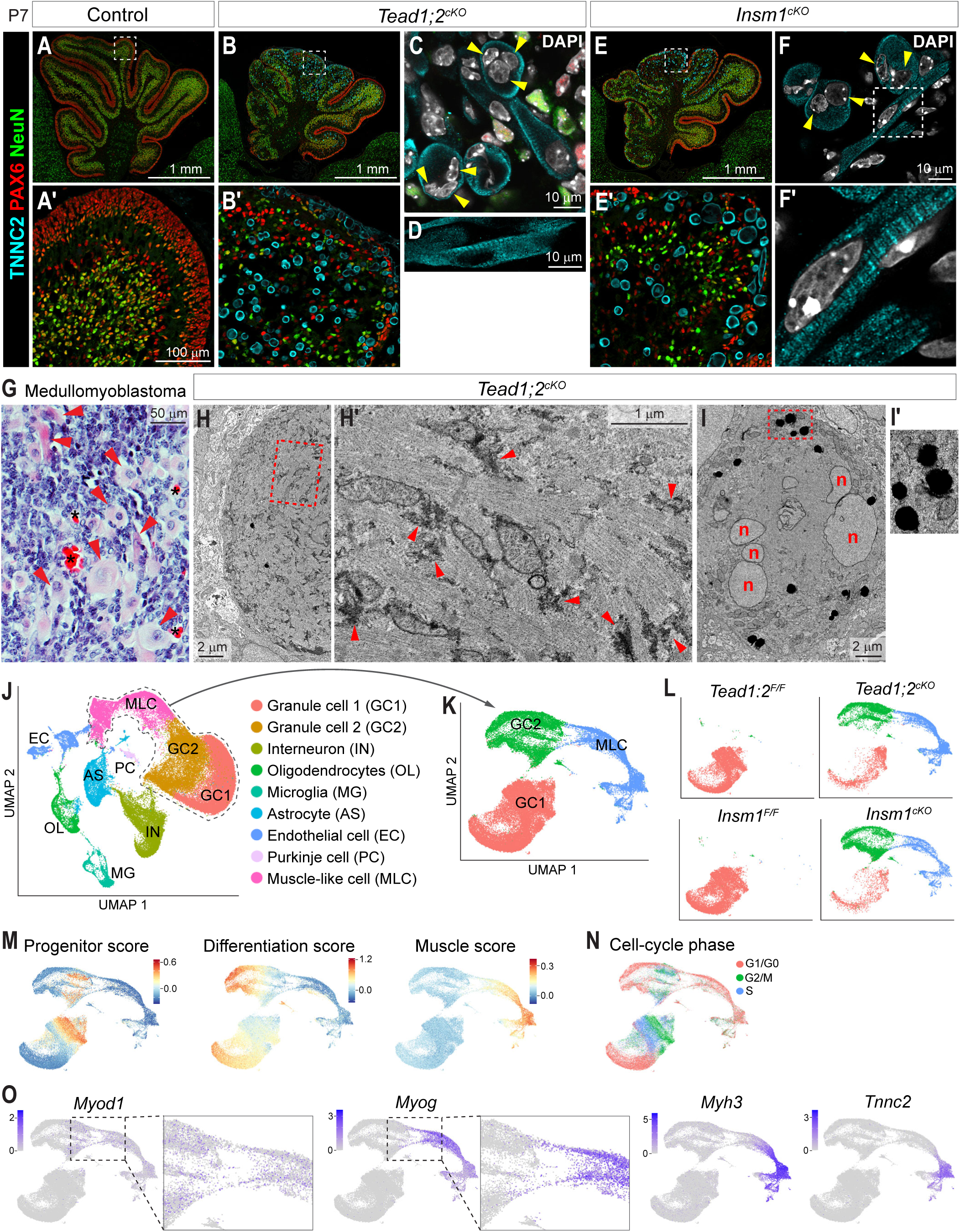
Characterization of muscle-like cells in *Tead1;2^cKO^* and *Insm1^cKO^* cerebella. (**A** to **F′**) Co-immunostaining of P7 cerebella for PAX6, NeuN, and the muscle structural protein troponin C2 (TNNC2). Single confocal *z* section showing multinucleated cells (C and F, arrowheads) and elongated cells containing TNNC2-labeled cytoskeletons with sarcomere-like striations (D and F′). (**G**) Hematoxylin and eosin-stained section of a medulloblastoma with myogenic differentiation, highlighting large rhabdomyoblasts (pink cells, arrowheads). Asterisks, red blood cells. (**H** and **H′**) Electron micrographs of a muscle-like cell densely packed with actin filaments, highlighting actin bundles and glycogen granules (arrowheads; H′). (**I** and **I′**) Electron micrographs of a muscle-like cell with multiple nuclei (n) and lipid droplets (I′). (**J**) UMAP embedding showing clustering and annotation of cells from P7 control and mutant cerebella (*N* = 73,789 cells after quality control). (**K** and **L**) Re-embedding of GC1, GC2, and MLC clusters (*N* = 45,696 cells; K) and separated by genotype (L): *Tead1;2^F/F^* (*n* = 2 mice), 9,727 cells; *Tead1;2^cKO^* (*n* = 3), 14,086 cells; *Insm1^F/F^* (*n* = 2), 10,953 cells; *Insm1^cKO^* (*n* = 3), 10,930 cells. (**M**) Signature scores for gene sets associated with GNPs (Progenitor), differentiating granule neurons (Differentiation), and skeletal muscle development (Muscle). (**N**) Inferred cell-cycle phases. (**O**) Feature plots of selected muscle genes. Boxed regions in A, B, E, F, H, and I are shown at higher magnification in A′, B′, E′, F′, H′ and I′.

The muscle protein–positive cells were often large and multinucleated (Fig. 4, B′, C, E′, and F, arrowheads). Some were elongated and contained TNNC2-labeled cytoskeletons with sarcomere-like striations (Fig. 4, D and F′). These muscle-like cells were never co-labeled with PAX6 or NeuN (Fig. 4, B′ and E′; fig. S4, D to L), indicating a complete loss of granule neuron lineage identity. Morphologically, they closely resembled rhabdomyoblasts observed in medullomyoblastoma (Fig. 4G, arrowheads), a variant of medulloblastoma characterized by myogenic differentiation (*36*); the origin of these rhabdomyoblasts has been elusive.

To further characterize these cells, we examined their ultrastructure by electron microscopy. Compared with normal neural cells, muscle-like cells in P7 mutant cerebella appeared darker, indicative of high cellular protein content (fig. S5, A and B). Their cytoplasm was densely packed with actin filaments, frequently organized into bundles resembling myofibrils (Fig. 4, H and H′; fig. S5, C and C′). In addition, these cells contained abundant glycogen granules (Fig. 4H′; fig. S5C′; arrowheads) and lipid droplets (Fig. 4, I and I′; fig. S5, D and D′), hallmarks of muscle cell metabolism. The presence of glycogen granules and lipid droplets was confirmed by Periodic Acid-Schiff and BODIPY staining, respectively (fig. S5, E to J′). Thus, loss of TEAD1/2 or INSM1 during cerebellum development induced a complete neural-to-muscle lineage conversion, with mutant cells acquiring not only muscle gene expression but also muscle-specific structural and metabolic features.

The abundance of muscle-like cells varied across cerebellar lobules. In both mutant lines, muscle-like cells were most abundant in lobules VI–VIII, where reductions in PAX6^+^ GNPs and NeuN^+^ granule neurons were also most pronounced (Fig. 4, B and E; fig. S4, A to C). However, far fewer TNNC2^+^ cells were present in anterior lobules (I–V), despite severe reductions in GNPs and granule neurons. Conversely, numerous TNNC2^+^ cells were present in lobule IX, where GNP and granule neuron populations were largely preserved. The lack of a simple correlation between muscle and neural phenotypes suggests that the two phenotypes arise through distinct mechanisms.

## Single-cell transcriptomic analysis reveals divergence of neural and muscle-like lineages

To comprehensively assess transcriptomic changes following TEAD1/2 or INSM1 loss, we performed single-cell RNA-seq (scRNA-seq) on P7 mutant cerebella (*n* = 3) and corresponding littermate controls (*n* = 2). Cell clusters were visualized by uniform manifold approximation and projection (UMAP) and annotated based on established marker genes (Fig. 4J). Two clusters—granule cell 2 (GC2) and muscle-like cell (MLC)—were composed almost entirely of mutant cells, whereas the granule cell 1 (GC1) cluster consisted predominantly of control cells (fig. S6A; also see Fig. 4L). The remaining clusters contained comparable proportions of control and mutant cells (fig. S6, B and C).

We next focused on the GC1, GC2, and MLC clusters. Re-embedding these cells by UMAP revealed clear separation between GC1 and GC2, with GC2 showing loose connectivity to MLC (Fig. 4K). Scoring cells using published granule neuron lineage progenitor and differentiation signatures (*37*) and our curated skeletal muscle development signatures (table S2) showed that GC1 and GC2 each contained both progenitor-like and differentiating granule cells (Fig. 4M), consistent with cell-cycle phase analysis (Fig. 4N). In contrast, MLC cells expressed muscle signatures but scored low for both progenitor and differentiation signatures (Fig. 4M), suggesting loss of granule neuron lineage identity. Whereas *Myod1* expression spanned GC2 and MLC clusters, *Myog* expression was largely restricted to MLC (Fig. 4O), suggesting that *Myog* upregulation was associated with departure from granule neuron lineage identity. Together, these results indicate that, in P7 mutant cerebella, the muscle-like population represents a lineage that has largely diverged from the neural lineage. Moreover, both progenitor-like and differentiating granule neuron populations in mutant cerebella were transcriptionally distinct from their normal counterparts.

## KDM1A interacts with TEAD and INSM1 and represses muscle fate during cerebellum development

The phenotypic and transcriptomic similarities between *Tead1;2^cKO^* and *Insm1^cKO^* cerebella suggest that TEAD1/2 act in concert with INSM1 during cerebellum development, promoting GNP proliferation and differentiation while simultaneously repressing muscle fate. INSM1 has been shown to interact with the transcriptional corepressor KDM1A (lysine demethylase 1A) through its SNAG domain (*38*). Immunostaining revealed that KDM1A was ubiquitously expressed during cerebellum development (fig. S7, A to G). To test whether TEAD, INSM1, and KDM1A form a tripartite complex, we performed co-immunoprecipitation experiments. In cerebellar nuclear extracts, immunoprecipitation of INSM1 or TEAD1 recovered KDM1A (Fig. 5, A and B). Moreover, in 293T cells—which do not endogenously express INSM1—immunoprecipitation of TEAD1 recovered KDM1A in the presence of transfected INSM1, but not in its absence or in the presence of INSM1^ΔSNAG^, a SNAG-domain deletion mutant exhibiting markedly reduced interaction with KDM1A but unperturbed interaction with TEAD1 (Fig. 5C). These results indicate that TEAD, INSM1, and KDM1A form a tripartite complex, with INSM1 serving as the bridging factor.

**Fig. 5.**
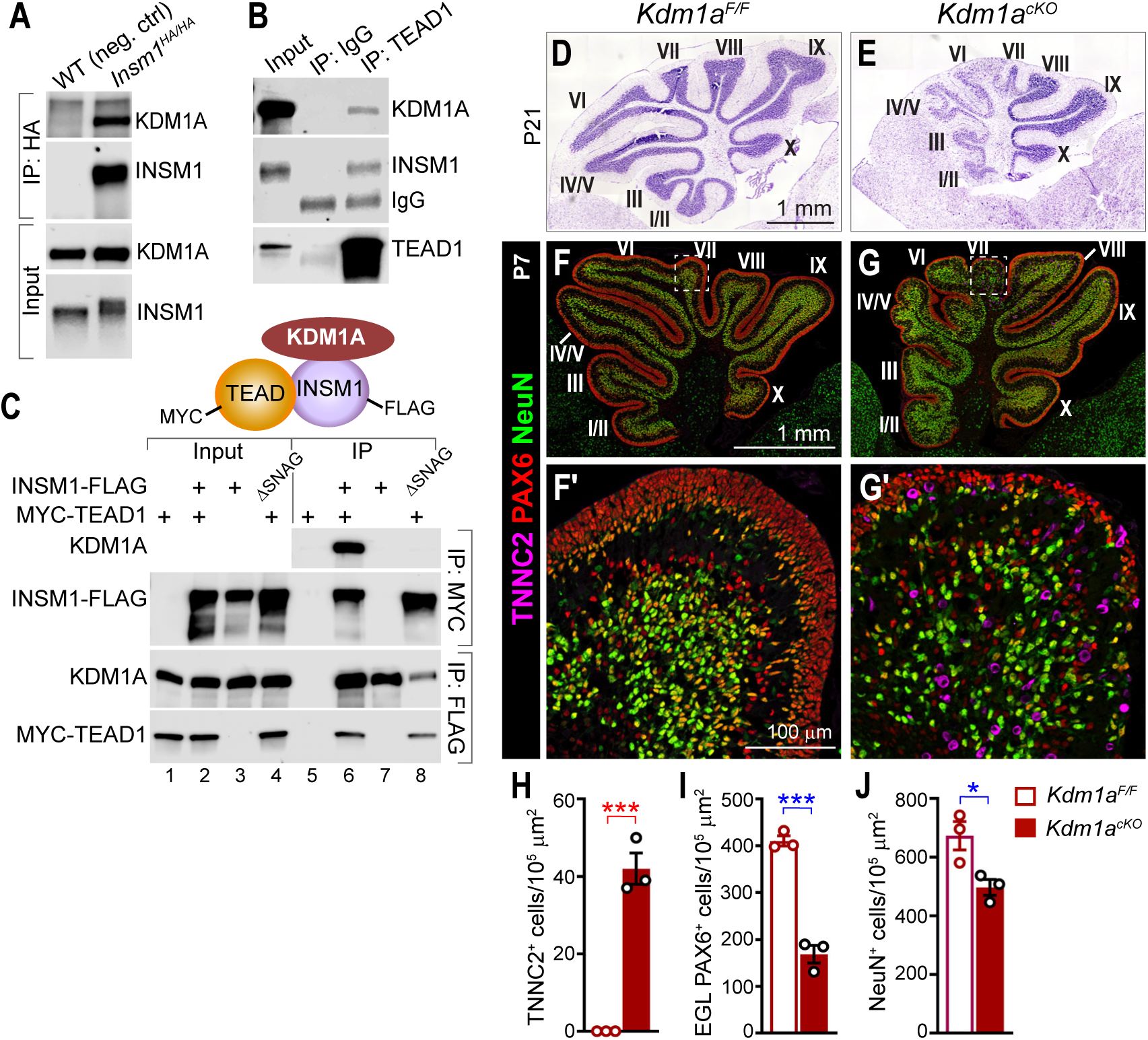
KDM1A interacts with TEAD and INSM1 and represses muscle fate during cerebellum development. (**A**) Co-immunoprecipitation (co-IP) of KDM1A by an anti-HA antibody from P7 *Insm1^HA/HA^*, but not WT (negative control), cerebellar nuclear extracts. (**B**) Co-IP from P7 WT cerebellar nuclear extracts using either a nonspecific IgG (negative control) or an anti-TEAD1 antibody. (**C**) Co-IP from 293T cells transfected with the indicated TEAD1 and INSM1 constructs. Endogenous KDM1A was detected. (**D** and **E**) Nissl staining of P21 cerebellum vermis sections from control and *Kdm1a^cKO^* mice. (**F** to **G′**) Co-immunostaining of P7 cerebella for PAX6, NeuN, and TNNC2. Boxed regions in F and G are shown at higher magnification in F′ and G′, respectively. (**H** to **J**) Quantifications of lobule VII TNNC2^+^ cells, PAX6^+^ cells located in the external granule layer (EGL), and NeuN^+^ cells. Each data point represents an individual animal. Values are mean ± SEM. Unpaired two-tailed *t*-test; *, *P* < 0.05; ***, *P* < 0.001.

To determine whether KDM1A is involved in repressing muscle fate during cerebellum development, we deleted *Kdm1a* using *Atoh1-Cre*. At P21, lobules I–VII of *Kdm1a^cKO^* cerebella were reduced in size and hypocellular compared with controls (Fig. 5, D and E). At P7, numerous TNNC2^+^ cells were present in *Kdm1a^cKO^* cerebella, predominantly in lobules VII–IX (Fig. 5, F to H; fig. S7, H and I), while PAX6^+^ GNPs and NeuN^+^ granule neurons were reduced (Fig. 5, I and J). These phenotypes resemble those of *Tead1;2^cKO^* and *Insm1^cKO^* cerebella, albeit with reduced severity, supporting a role for KDM1A in TEAD–INSM1-mediated repression of muscle fate during cerebellum development.

## Loss of TEAD1/2 or INSM1 reshapes chromatin accessibility in GNPs

To investigate the gene regulatory mechanisms by which TEAD1/2 and INSM1 regulate cerebellum development, particularly their role in repressing muscle fate, we compared chromatin accessibility in P2 mutant and littermate control cerebella using ATAC-seq and profiled genome-wide TEAD1 and INSM1 chromatin-binding by CUT&RUN in P2 WT (for TEAD1) or *Insm1^HA/HA^* (for INSM1) cerebella (fig. S8A). Even at this early stage, loss of TEAD1/2 or INSM1 caused widespread changes in chromatin accessibility, with 34,665 and 70,154 regions significantly altered (false discovery rate < 0.05) in *Tead1;2^cKO^* and *Insm1^cKO^* cerebella, respectively, corresponding to 19% and 32% of all accessible regions detected in each mouse line (Fig. 6A; fig. S8B). Genomic Regions Enrichment of Annotations Tool (GREAT) analysis revealed significant enrichment of muscle development–related gene sets among the regions that gained accessibility, whereas the regions that lost accessibility were enriched for gene sets associated with cerebellum or neural development (table S3).

**Fig. 6.**
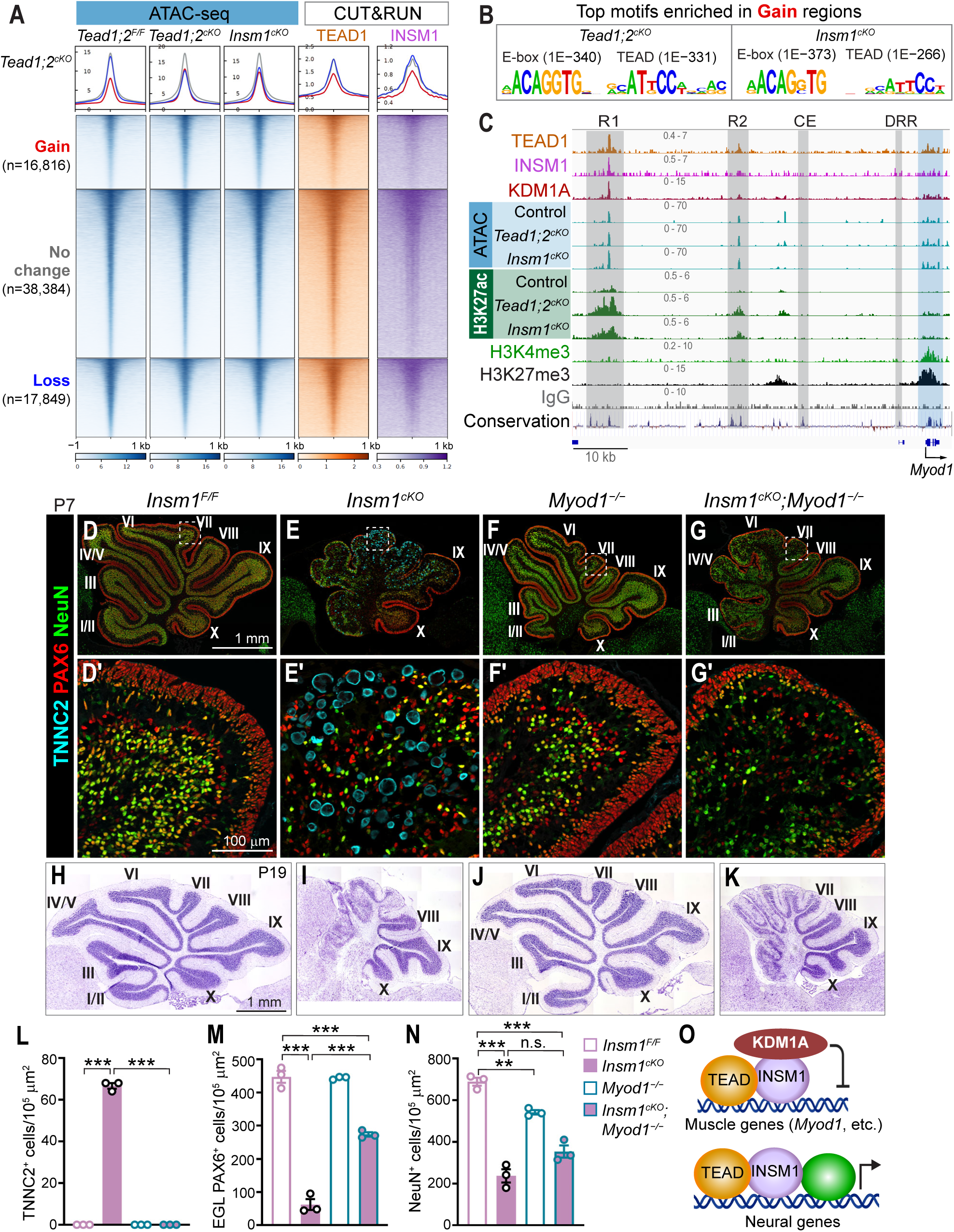
*Myod1* upregulation is required for muscle lineage conversion following INSM1 loss. (**A**) Profile graphs and heatmaps of chromatin accessibility (ATAC-seq) in P2 control and mutant cerebella, and TEAD1 and INSM1 genomic binding (CUT&RUN) in P2 WT (*Insm1^HA/HA^* for INSM1) cerebella, grouped by their change in accessibility in *Tead1;2^cKO^* cerebella compared with littermate controls. Gain and Loss, FDR < 0.05, no fold-change (FC) cutoff; No change, |FC| < 1.05, *P* > 0.05. Profile graphs and heatmaps of ATAC-seq data were scaled based on the signals of no-change regions. One representative replicate of 2–4 biological replicates. (**B**) Top enriched motifs in the regions with gain of accessibility in mutant cerebella compared with no-change regions, with *P* values. (**C**) The *Myod1* locus showing transcription factor binding, chromatin accessibility (ATAC), and histone marks in P2 cerebella. One representative replicate of 2–4 biological replicates. Conservation profile (100 vertebrates) was obtained from USCS genome browser. (**D** to **G′**) Co-immunostaining of P7 cerebella for PAX6, NeuN, and TNNC2. Boxed regions in D to G are shown at higher magnification in D′ to G′. (**H** to **K**) Nissl staining of P19 cerebellum vermis sections from control and mutant mice. (**L** to **N**) Quantifications of lobule VII TNNC2^+^ cells, PAX6^+^ cells located in the external granular layer (EGL), and NeuN^+^ cells. Each data point represents an individual animal. Values are mean ± SEM. Unpaired two-tailed *t*-test; **, *P* < 0.01; ***, *P* < 0.001; n.s., not significant (*P* > 0.05). (**O**) Model depicting dual roles of TEAD and INSM1 in repressing muscle genes and facilitating neural gene expression during cerebellum development.

The regions that gained accessibility in mutant cerebella exhibited, on average, lower accessibility and reduced TEAD1/INSM1 binding in WT cerebella than either unchanged regions or regions that lost accessibility (Fig. 6A; fig. S8B). These regions were highly enriched for the CAGNTG E-box motif preferred by MRFs (*39*), followed by the TEAD motif (Fig. 6B; fig. S8C). Restricting the analysis to promoter-distal, putative enhancer regions yielded similar results (fig. S8, D to F). Together, these findings suggest that many regions gaining accessibility are not repressed by direct TEAD/INSM1-binding and that loss of TEAD1/2 or INSM1 renders the chromatin landscape more permissive to engagement by bHLH TFs preferring the CAGNTG E-box, such as MRFs.

We next asked whether derepression or activation of myogenic master regulators contributed to reshaping the chromatin landscape. Two MRFs, *Myod1* and *Myog*, were upregulated following TEAD1/2 or INSM1 loss (Fig. 3B; fig. S3A; table S1). Given that MYOD functions upstream of MYOG (*33*) and is capable of reprogramming diverse cell types to the skeletal muscle lineage (*40–42*), we focused on *Myod1* regulation. In P2 WT cerebellum, the *Myod1* locus was accessible and poised, harboring both the active promoter mark H3K4me3 and the repressive mark H3K27me3 (Fig. 6C, blue-shaded region). Furthermore, two upstream regions were accessible and bound by TEAD1, INSM1, and KDM1A in WT cerebellum and gained the active enhancer mark H3K27ac in *Tead1;2^cKO^* and *Insm1^cKO^* cerebella (Fig. 6C, R1 and R2), suggesting that these elements contribute to *Myod1* upregulation in mutant cerebella. Both regions are highly conserved in vertebrates (Fig. 6C).

Notably, the two enhancers that regulate *Myod1* during normal skeletal muscle development, the core enhancer (CE) and the distal regulatory region (DRR) (*43*), were inaccessible in both WT and mutant cerebella and did not acquire H3K27ac following TEAD1/2 or INSM1 loss (Fig. 6C), indicating that they are not involved in *Myod1* upregulation in mutant cerebella. Thus, *Myod1* expression is controlled by distinct *cis*-regulatory elements during cerebellum and skeletal muscle development.

## MYOD upregulation following INSM1 loss drives muscle lineage conversion

To test whether *Myod1* upregulation is required for muscle lineage conversion, we deleted *Myod1* in the *Insm1^cKO^* background. We did not attempt *Myod1* deletion in the *Tead1;2^cKO^* background because all three genes are located on the same chromosome, making triple deletion technically challenging. Loss of *Myod1* during normal development does not prevent skeletal muscle formation owing to compensation by *Myf5* (*44*). *Myod1* loss also did not overtly disrupt cerebellum development. At P7, *Myod1^−/−^* cerebella displayed overall normal morphology, albeit with reduced size compared with controls (Fig. 6, D and F), likely associated with the smaller body size of the animals (fig. S9). By P19, *Myod1^−/−^* cerebella appeared grossly normal (Fig. 6J).

Strikingly, loss of *Myod1* completely prevented the appearance of muscle-like cells in the *Insm1^cKO^* background (Fig. 6, D to G′, and L), demonstrating that MYOD is required for muscle lineage conversion following *Insm1* loss during cerebellum development.

At P7, GNPs and granule neurons remained significantly reduced in *Insm1^cKO^;Myod1^−/−^* double-mutant cerebella, although GNP numbers were partially restored relative to *Insm1^cKO^* cerebella (Fig. 6, D to G′, M, and N). Accordingly, at P19, lobules I–VII of double-mutant cerebella remained severely malformed, albeit with improved morphology compared with *Insm1^cKO^* cerebella. These findings indicate that neural proliferation and differentiation defects are only partially attributable to MYOD upregulation, supporting the conclusion that repression of muscle fate and promotion of neural development represent separable functions of INSM1 during cerebellum development (Fig. 6O).

## Discussion

Cell fate commitment during development is commonly viewed as involving progressive restriction of developmental potential, enforced largely by passive, heterochromatin-based silencing of alternative lineage programs (*6, 7*). Our findings challenge and expand this view by showing that the maintenance of neural lineage identity in the developing cerebellum requires active repression of a vastly divergent fate. Specifically, we show that the TEAD–INSM1 transcriptional complex is essential not only for promoting GNP proliferation and differentiation, but also for actively repressing skeletal muscle identity (Fig. 6O). Loss of this repression upregulates the myogenic master regulator *Myod1* and drives complete neural-to-muscle lineage conversion in vivo.

*Myod1* upregulation likely requires not only release from TEAD–INSM1-mediated repression, but also activation by a transcriptional activator, whose identity remains unknown. Notably, rare MYOD^+^ and MYOG^+^ cells are detectable even in WT cerebella. These cells progressively lose granule neuron lineage identity but do not undergo full muscle differentiation, suggesting that TEAD–INSM1 also represses the myogenic differentiation program downstream of MYOD/MYOG. Whether these rare cells represent stochastic breaches of lineage fidelity during normal development or serve a physiological role remains to be determined.

Our findings also raise the question of whether cerebellar GNPs are uniquely susceptible to this fate conversion. Loss of TEAD1/2 or INSM1 in multiple regions of the developing nervous system—including the telencephalon (*20, 45, 46*), inner ear (*47*), and retina (*48*)—does not elicit muscle conversion, although ectopic expression of muscle structural genes have been reported in the *Insm1*^−/−^ pituitary gland (*38*). The myogenic program may be more stably silenced in regions that resist conversion. Alternatively, these regions may lack the transcriptional activity required to activate *Myod1*, or the neural identity of GNPs may be intrinsically more labile. Whether other neural cell types are susceptible to muscle conversion remains to be determined, as does whether the greater susceptibility of GNPs is linked to their intrinsic features such as their exceptional proliferative capacity or prolonged developmental window.

Although the phenotypically complete, cross–germ-layer lineage conversion we see here is striking, we interpret it not as evidence of broadly heightened plasticity in cerebellar GNPs, but rather as indicating a selective vulnerability of the neural–muscle fate boundary relative to other lineage barriers. Supporting this idea, disruption of Polycomb-mediated repression during cerebellum development aberrantly upregulates muscle genes without activating other lineage programs (*49*). Moreover, TEAD depletion in primary myoblasts induces neural gene expression while impairing muscle differentiation (*50*). Mixed-lineage tumors further underscore this bidirectional instability: medullomyoblastomas exhibit myogenic differentiation within neural tumors (*36*), rare pineoblastomas show rhabdomyoblastic components (*51*), and subsets of rhabdomyosarcomas contain cells expressing neural genes (*52*). Even under normal conditions, muscle fibers have been observed in the pineal glands of rats and pigs (*53, 54*).

An evolutionary perspective may help explain this selective vulnerability. Neural and muscle lineages are thought to be sister branches derived from a common ancestral cell type, such as myoepithelial cells (*55–57*), and share deeply conserved regulatory architectures, including cascades of bHLH TFs. This evolutionary proximity may help explain why at least a subset of neural progenitors retain latent competence to activate myogenic programs. We propose that integrating developmental and evolutionary perspectives on cell fate will be particularly informative for understanding how lineage boundaries are enforced—and breached—during normal development and disease.

## Materials and Methods

### Mice

Mice were maintained with a standard 12-h light/dark schedule. Both male and female mice were used for experiments. All animal procedures were approved by the Institutional Animal Care and Use Committee of St. Jude Children’s Research Hospital (SJCRH).

The *Atoh1-Cre* (*30*) (strain #011104), *Kdm1a^F/F^* (*58*) (#023969), and *Myod1^−/−^* (*59*) (#002523) lines were obtained from the Jackson Laboratory. The *Insm1^F/F^* (*47*) and *Tead2^F/F^* (*60*) lines were provided by Jaime García-Añoveros (Northwestern Feinberg School of Medicine) and Melvin DePamphilis (National Institutes of Health), respectively. Generation of *Tead1^F^* and *Tead1;2^F^* alleles was described previously (*20*).

The *Insm1^HA^* knock-in mouse allele was generated by coinjecting guide RNA targeting *Insm1* locus (5′-GACCGAGGGCGCACTCTAGCNGG-3′), donor ssDNA (sense strand megamer, Integrated DNA Technologies [IDT]) encoding 2 copies of HA-tag and one copy of AM (Active Motif) tag, and Cas9 protein into the pronucleus of one-cell-stage C57BL/6J zygotes. F0 mice were screened for targeted integration by junction PCR and sequencing, and those with correct integration were crossed with WT mice for germline transmission. F1 mice were further validated using junction PCR and sequencing. The following primers were used to genotype the WT and knock-in alleles: F, 5′-TAGACAGGTGATCCTCCTTCAG-3′; R, 5′-TTCACCCTGGAATGGGAGTA-3′, with the WT allele being ∼190 bp and the knock-in allele ∼310 bp.

For cell-cycle exit analysis, EdU (5-ethynyl-2′-deoxyuridine) was injected into postnatal day 2 (P2) mice intraperitoneally at 10 μg per gram body weight. Animals were collected 24 h later.

### Tissue collection, histology, and immunostaining

Embryonic mouse brains were dissected in cold PBS and fixed overnight in freshly prepared 4% paraformaldehyde (PFA; Electron Microscopy Sciences [EMS] #15710) at 4°C. Postnatal animals were transcardially perfused with PBS and PFA, and their brains were dissected and fixed in PFA overnight at 4°C. For frozen sections, fixed brains were washed in PBS, equilibrated in 30% sucrose overnight at 4°C, embedded in O.C.T. Compound (SAKURA #4583), and sectioned at 12–14-μm thickness. Nissl staining, hematoxylin and eosin (H&E) staining, and periodic acid-Schiff (PAS) staining were performed according to standard protocols.

For immunofluorescence staining, frozen sections were washed in PBS, blocked and permeabilized in PBS with 0.5% Triton X-100 (PBST) and 10% normal donkey serum (Jackson Immuno Research #017-000-121) for 1 h, and incubated in primary antibodies diluted in PBS supplemented with 0.1% Triton X-100 and 5% normal donkey serum at 4°C overnight. Sections were then washed in PBST and incubated in fluorescence–conjugated secondary antibodies (Jackson Immuno Research) at 1:1000 dilution and DAPI (1:1000 of 1 mg/ml stock; Sigma #32670) for 2–3 h at room temperature. Sections were then washed in PBST and mounted in ProLong Gold antifade mountant (Invitrogen #P36931). The following primary antibodies were used: TEAD1 (Cell Signaling Technology [CST] #12292, 1:500), phospho-Histone H3 (Ser10) (Millipore Sigma #06-570, 1:500), HA-Tag (CST #3724, 1:500), PAX6 (R&D Systems #AF8150-SP, 1:300), Ki67 (Abcam ab15580, 1:500), BLBP (R&D Systems #AF3166, 1:500), MYOG (Abcam #ab124800, 1:500), MYOD (Novus Biologicals #NBP2-32882, 1:50), NeuN (Synaptic Systems #266004, 1:300), TAG1 (DSHB #4D7, 1:50), Calbindin (Synaptic Systems #214005, 1:500), KDM1A (Abcam #ab17721, 1:500), MYH (DSHB #MF20-c, 1:50), MYH I (DSHB #BA-F8, 1:50), MYH IIA (DSHB #SC-71, 1:50), MYH IIB (DSHB #BF-F3, 1:50). MYL1 (ProteinTech #15814-1-AP, 1:250), TNNC2 (ProteinTech #15875-1-AP, 1:250), MYLPF (ProteinTech #16052-1-AP, 1:250), and Cav3 (ProteinTech #28358-1-AP, 1:250). EdU staining was performed using the Click-iT Plus EdU kit (Thermo Fisher Scientific #C10638) according to the manufacturer’s protocol. TUNEL assay was performed using the ApopTag Fluorescein *In Situ* Apoptosis Detection kit (Millipore Sigma #S7110) according to the manufacturer’s protocol. BODIPY (Invitrogen #D3922, 1:1000) staining was performed according to the manufacturer’s protocol.

### Image acquisition and quantification

Bright field images were acquired using a Zeiss AxioImager with a 10× objective lens. Fluorescence images were acquired using a Zeiss LSM780 confocal microscope or a Zeiss Apotome microscope with a 20× or 40× objective lens, or a Zeiss LSM 980 AiryScan microscope with 60× objective lens. For quantification of PAX6^+^ and NeuN^+^ cells and cell-cycle exit index, regions of interest (ROIs) were first outlined and extracted in Zeiss Zen 3.11 software, nucleus segmentation was then performed using the machine-learning–based segmentation method StarDist (https://github.com/stardist/stardist), which was first trained using manually annotated datasets. Fluorescent label intensity of each channel over nuclear ROIs was calculated using a custom multi-channel intensity quantification algorithm created in OMERO (https://www.openmicroscopy.org/omero/scientists/); thresholds for signal positivity were manually determined by inspecting each image. MYOD^+^, MYOG^+^, and TNNC2^+^ cells were counted manually. TAG1 signal intensity and tissue area were measured using Fiji software. Data points in all quantification graphs correspond to individual animals, with each point representing the average of 3 sections per animal.

### Electron microscopy

Postnatal animals were transcardially perfused with PBS and freshly prepared 2.5% glutaraldehyde/2% PFA in 0.5 M phosphate buffer, and their brains were dissected and fixed in 2.5% glutaraldehyde/2% PFA in 0.5 M phosphate buffer overnight at 4°C. Vibratome sections of 100-µm thickness were prepared for electron microscopy as described (*61*). Briefly, tissue sections were rinsed in 0.1 M cacodylate buffer (pH7.4) for 3× 5 min before incubating for 90 min in the same buffer containing 40 mM osmium tetroxide, 35 mM potassium ferrocyanide, and 2.5 M formamide (Sigma Aldrich) followed, without washing, with incubation in 40 mM osmium tetroxide in 0.1 M cacodylate buffer for 90 min. Sections were then washed in 0.1 M cacodylate buffer for 3× 5 min, incubated in 320 mM pyrogallol (Sigma Aldrich) in water for 30 min, washed in water for 3× 5 min, and incubated in 40 mM osmium tetroxide in water for 90 min. Following washing in water for 3× 5 min, samples were incubated in Walton’s lead aspartate solution (*62*) at 60°C for 1 h, washed in water for 3× 5 min, dehydrated in an ascending ethanol series (30% to 100%) on ice, transitioned in propylene oxide, and infiltrated with EmBed812 resin. Resin was polymerized at 60°C for 48 h. Ultrathin sections of 80-nm thickness were cut on a Leica UC-7 ultramicrotome with a Diatome diamond knife and mounted either on copper transmission electron microscope (TEM) grids or silicon chips. Sections on TEM grids were scanned using a Tecnai F20 TEM (Thermo Fisher) operating at 80 kV and equipped with a NanoSprint15 MkII camera (Advanced Microscopy Techniques). Sections on silicon chips were mounted in Leitsilber 200 silver paint (Ted Pella #16035) onto standard ∅ 12.7 mm pin stubs (Ted Pella #16111-9) and scanned under a Zeiss GeminiSEM 460 using a Sense backscattered electron detector. Unless otherwise noted, all reagents were obtained from EMS.

### Plasmids

DNA fragments encoding murine INSM1 (ID: 53626) with 2× FLAG at the C-terminus and murine TEAD1 (ID: 21676) with an N-terminal MYC-tag were synthesized by Bio Basic USA or IDT (gBlock) and cloned into pcDNA3.1(+) vector by Gibson assembly using NEBuider HiFi DNA Assembly Master Mix (New England Biolabs [NEB] #E2621).

### Cell culture and transfection

293T cells were obtained from ATCC (CRL-3216) and maintained in DMEM with high glucose supplemented with 10% FBS and penicillin/streptomycin. Four μg of DNA mixture was transfected into 6 × 10^6^ 293T cells cultured in a 6-cm dish with 3.5 ml of medium without antibiotics, using 12 µl of Lipofectamine 2000 (Thermo Fisher Invitrogen #11668027) according to the manufacturer’s instruction. Cells were cultured for 24 h, repropagated into a 10-cm dish with 12 ml of medium, and cultured for another 24 h before being collected for co-immunoprecipitation experiments.

### Co-immunoprecipitation

P7 mouse cerebella were dissected in PBS, flash-frozen in dry ice-ethanol slurry, and stored at −80°C. One cerebellum was used for each co-immunoprecipitation (co-IP) reaction. Each 10-cm plate of transfected 293T cells were used for two co-IP reactions. Co-IP was performed using Nuclear Complex Co-IP kit (Active Motif #54001) according to kit instruction, using IP Low buffer, and 15 µl of TEAD1 (CST #12292, 7 ng/µl), 3 µl of HA (Abcam #ab9110, 1 µg/µl), 3 µl of FLAG-M2 (Sigma #F1804, 1 µg/µl), or 15 µl of MYC-9E10 (Santa Cruz sc-40, 0.2 µg/µl) antibody. The following primary antibodies were used for Western blotting: TEAD1 (CST #12292, 1:500), INSM1 (Santa Cruz sc-271408, 1:500), KDM1A (CST #2139, 1:1000), FLAG (CST #2368S, 1:1000), and MYC (CST #2278, 1:500).

### Bulk RNA-seq and data analysis

P2 cerebella were dissected in cold PBS. RNA was isolated using the Direct-zol RNA MiniPrep kit (Zymo Research #R2052) according to the manufacturer’s protocol and quantified using the Quant-iT RiboGreen RNA assay (Thermo Fisher #R11491) and quality checked by the 2100 Bioanalyzer RNA 6000 Nano assay (Agilent) or 4200 TapeStation High Sensitivity RNA ScreenTape assay (Agilent). Sequencing libraries were prepared using the Illumina Stranded Total RNA Prep library kit (Illumina #20040529) according to the manufacturer’s instructions, analyzed for size distribution using the 2100 BioAnalyzer High Sensitivity kit (Agilent), 4200 TapeStation D1000 ScreenTape assay (Agilent), or 5300 Fragment Analyzer NGS fragment kit (Agilent), and quantified using the Quant-iT PicoGreen ds DNA assay (Thermo Fisher) or by low pass sequencing with a MiSeq nano kit (Illumina). Paired end 100 cycle sequencing was performed on a NovaSeq 6000 sequencer (Illumina).

Raw reads were trimmed using TrimGalore v0.6.3 (*63*), then mapped to the *Mus musculus* reference genome (GRCm38.p6) using STAR v2.7.9a (*64*) in a splice-site aware manner based on Gencode M22 primary assembly annotations. Gene level quantification was determined using RSEM v1.3.3 (*65*). Read counts were analyzed in R v4.1.0 using the limma-voom pipeline (*66*). The analysis was restricted to confidently annotated (levels 1 and 2) protein-coding genes. Low-count genes were filtered out using a CPM (counts per million) cutoff of 0.1. To account for compositional biases, normalization factors were calculated using the Trimmed Mean of M-values (TMM) method. The counts were then transformed via the *voom* function to estimate the mean-variance relationship and assign precision weights to each observation. Differential expression was modeled using the *lmFit* and *eBayes* functions within the limma v3.42.2 package.

### Single-cell RNA-seq and data analysis

P7 mouse cerebella were dissected in PBS and dissociated individually into single-cell suspensions using a papain-based Neural Tissue Dissociation Kit (Miltenyl Biotec #130-094-802) according to the manufacturer’s protocol. Cells were filtered through a 40-μm strainer (pluriSelect 43-10040-40), spun down at 800 × *g* for 3 min, resuspended in PBS, and counted on a Countess automated cell counter (Invitrogen). One million cells per animal were fixed using the Parse Biosciences Evercode Cell Fixation v3 kit (ECFC3300) according to the manufacturer’s protocol, slow-frozen in a Mr. Frosty Freezing container (Thermo Fisher #5100-0001), and stored at −80°C in aliquots for up to 2 months. Sequencing libraries were generated using the Parse Evercode™ WT kit (ECWT3300) according to the manufacturer’s protocol, combining 3 *Tead1;2^cKO^*, 2 *Tead1;2^F/F^*; 3 *Insm1^cKO^*, and 2 *Insm1^F/F^* samples. Libraries were analyzed for size distribution on an Agilent TapeStation and sequenced on an Illumina NovaSeq X Plus sequencer.

FASTQ files were aligned to the mouse reference genome (mm10/GRCm38) and processed in the Parse Trailmaker cloud suite. Cells with ≥800 UMIs and <5% mitochondrial gene content were retained. Doublets were identified and removed using Trailmaker’s built-in doublet detection algorithm. The resulting Seurat object (73,789 cells across 10 samples) was downloaded for local analysis in R (Seurat v4.4.0). Ensembl gene IDs were converted to gene symbols using the annotation table stored in the misc slot of the Parse-exported Seurat object; Ensembl IDs were retained for genes lacking a corresponding symbol. Data were log-normalized using *NormalizeData* with a scale factor of 10,000.

The 2,000 most variable genes were identified by variance-stabilizing transformation (*FindVariableFeatures*, method = "vst"). Data were scaled and PCA was performed on the variable features. UMAP was computed based on the top 20 PCs. A shared nearest-neighbor graph (*FindNeighbors*, dims = 1:20) was constructed and Leiden clustering applied (*FindClusters*, algorithm = 4, method = "igraph", resolution = 0.1, random.seed = 1234). No batch integration was applied. Cell types were annotated using established markers for granule cells (*Atoh1*, *Insm1*, *Barhl1*), interneurons (*Pax2*, *Gad1*), astrocytes (*Aldoc*, *Aqp4*), microglia, (*P2ry12*, *Sall1*), oligodendrocytes (*Olig1*, *Olig2*), Purkinje cells (*Calb1*, *Pcp2*), endothelial cells (*Pecam1*, *Cdh5*), and muscle-like cells (*Myog*, *Myod1*, *Myh3*). The statistical significance of differences in cell type proportions was tested using propeller (*67*). Cell cycle phase was scored using Seurat’s *CellCycleScoring* function using the cc.genes.updated.2019 S and G2/M gene signatures, converted to title case for mouse gene nomenclature compatibility.

Granule cell and muscle-like cell clusters (GC1, GC2, and MLC) were subset, and cells with fewer than 750 detected features were excluded. The subset was reclustered using the top 3,000 variable features and 50 PCs (UMAP spread = 1.5, min.dist = 0.1). Granule cell progenitor and differentiation signature genes were obtained from published gene lists (*37*). Muscle signature genes were compiled by combining genes in the following GOBP gene sets: GO:0035914 (skeletal muscle cell differentiation), GO:0014856 (skeletal muscle cell proliferation), GO:0048741 (skeletal muscle fiber development), GO:0098528 (skeletal muscle fiber differentiation), and GO:0060538 (skeletal muscle organ development), then removing genes also present in granule cell signature gene lists (*37*) (10 genes were removed), yielding 227 genes. Gene signature scores were calculated using the *AddModuleScore* function.

### ATAC-seq and data analysis

P2 mouse cerebella were dissected and dissociated using a Neural dissociation kit (Miltenyl Biotec 130-094-802) according to the manufacturer’s protocol. Briefly, tissues were incubated in a papain-based solution for 10 min at 37°C with occasional mixing, dissociated by pipetting, filtered through a 5-ml tube with cell-strainer cap (Falcon 352235), and spun down at 800 × *g* for 3 min. Cells were resuspended in PBS, counted, and 100,000 cells were aliquoted into 200 µl of BAMBANKER cell freezing medium (Bulldog Bio Inc BB05) and stored at −80°C. ATAC-Seq reactions were performed using an ATAC-seq kit (Active Motif 53150) according to the manual. Libraries were checked on Agilent 4200 TapeStation using D1000 ScreenTape assay for size distribution, then pooled and sequenced on an Illumina NextSeq 2000 or NovaSeq 6000 sequencer. For the *Insm1* line, 3 control and 3 mutant cerebella were analyzed. For the *Tead* line, 3 control and 4 mutant cerebella were analyzed.

Raw reads in fastq format were processed with Trim-Galore v0.4.4 (*63*) and cutadapt (DOI:10.14806/ej.17.1.200) to remove adaptors, followed by FastQC analysis. A quality score cutoff of Q20 was used. The first 15-bp of each read were also removed to reduce the GC bias. Next, reads were mapped to the mouse reference genome (mm10; GRCm38.p6) with BWA mem 0.7.17-r1188 (*68*), then converted to BAM format and deduplicated with fq2bam v3.0.0.6. Uniquely mapped, properly paired reads were retained using samtools v1.2 (*69*) and fragments were extracted using bedtools v2.24.0 (*70*). Fragments <2,000-bp were extracted and the center 80-bp of each fragment were used to generate bigwig track file using UCSC tools v4 (*71*). All tracks were normalized to 20 million fragments. Genome browser tracks were plotted using IGV v2.18.1 (*72*).

Peaks were first called for each sample using MACS2 (v2.1.1.20160309) (*73*). For each genotype, a reproducible peak was defined as a peak that was called with high confidence (FDR < 0.05) in at least one biological replicate and was also called as a peak (FDR < 0.5) in all the other replicates. Reproducible peaks within each genotype were then concatenated, sorted, and merged to generate genotype-specific reproducible peak sets.

To identify differentially accessible regions, a reference peak set was first generated by combining all reproducible peaks from mutant and control samples. Counts of reads in each sample overlapping the reference peak set were generated using the *intersect* command from pybedtools v0.9.1 (*70, 74*), and counts per peak were then converted to FPKM (fragments per kilobase per million mapped reads) unit for visualization and summary purposes. Batch effect correction was performed using ComBat-seq (v3.54.0) (*75*). TMM from edgeR (*76*) followed by limma-voom approach (*66, 77*) was used to assess the significance of differential accessibility. No-change regions were defined as |FC| < 1.05, *P* > 0.5, and average counts in both mutant and control groups > 0. Peaks with a summit >2,000-bp from a transcription start site (TSS), based on the reference gene annotation from Gencode vM14 (*78*), were considered promoter-distal.

GREAT (Genomic Regions Enrichment of Annotations Tool) analysis (*79, 80*) was performed using default parameters with the whole reference peak set as the background. Motif enrichment analysis was performed using HOMER v4.10 (*81*) with no-change regions as the background.

### CUT&RUN and data analysis

P2 mouse cerebella were dissected and dissociated using a Neural dissociation kit (Miltenyl Biotec 130-094-802) according to the manufacturer’s protocol. Briefly, tissues were incubated in a papain-based solution for 10 min at 37°C with occasional mixing, dissociated by pipetting, filtered through a 5-ml tube with cell-strainer cap (Falcon 352235), and spun down at 800 × *g* for 3 min. Cells were resuspended in PBS, counted, spun down again, and resuspended in BAMBANKER cell freezing medium (Bulldog Bio Inc BB05) at 10^6^ cells per 200 µl and stored at −80°C.

One-million (for histone marks) or two-million (for transcription factors) cells were used for each CUT&RUN reaction, which was performed based on the protocol developed by the Henikoff lab (*82*) with minor modifications: nuclei were prepared according to EpiCypher CUTANA CUT&RUN protocol (v1.7); antibody, pAG-MNase, and CaCl_2_ incubations were carried out at room temperature for 4 h, 15 min, and 15 min, respectively; DNA was purified using CUTANA DNA purification column (EpiCypher #14-0050) or Monarch Spin PCR & DNA Cleanup Kit (NEB #T4130). For H3K27ac, cells were first cross-linked in 1% PFA at room temperature for 2 min and stopped in 125 mM glycine for 5 min before nuclei isolation, and released DNA fragments were treated with 0.1% SDS and 0.25 mg/ml proteinase K (Ambion #AM2546) at 65°C for at least 2 h before purification. The following antibodies per reaction (100 µl) were used: TEAD1 (CST #12292, 1 µl), HA (Abcam #ab9110, 1 µl), KDM1A (CST #2139, 5 µl), H3K27ac (Abcam #ab4729, 0.5 µl), H3K4me3 (Millipore Sigma #07-473, 1 µl), H3K27me3 (Active Motif #39155, 1 µl), and normal rabbit IgG (CST #2729).

Libraries were prepared using the NEBNext Ultra II kit (NEB #E7103) with modifications as in https://dx.doi.org/10.17504/protocols.io.bagaibse. Purified libraries were analyzed with TapeStation (Agilent), using the High Sensitivity D1000 reagents (Agilent 5067-5585), and sequenced on NovaSeq6000 sequencer to generate 50-bp paired-end reads.

Raw reads in fastq format were processed with Trim-Galore v0.4.4 and cutadapt (DOI:10.14806/ej.17.1.200) to remove adaptors, followed by FastQC analysis. A quality score cutoff of Q20 was used. Next, reads were mapped to the mouse reference genome (mm10; GRCm38.p6) with bwa aln, followed by bwa samse v0.7.12-r1039 (*68*) with -K flag set to 10,000,000, and the output was converted to BAM format with samtools v1.2 (*69*). Duplicated reads were marked with bamsormadup tool from biobambam2 v2.0.87 (DOI: 10.1186/1751-0473-9-13). Uniquely mapped, properly paired reads were extracted with samtools v1.2 (*69*) and sorted by reads name using biobambam2 v2.0.87. Bedtools v2.17.0 (*70*) were then used to convert bam files to bedpe files. Fragments <2,000 bp were kept for peak calling. MACS2 (v2.1.1.20160309) (*73*) and SICER (v1.1) (*83*) were used to identify sharp (transcription factors) and broad (histone marks) peaks, respectively. Fragments <2,000-bp were extracted and the center 80-bp of each fragment were used to generate bigwig track file by UCSC tools v4 (*71*). All tracks were normalized to 10 million fragments.

## Acknowledgments

We thank members of the Cao lab, especially Sandy Nguyen, and Alfonso Lavado and Yong Cheng for suggestions and technical help; the former lab member Ram Kumar for initiating the project; the Cell and Tissue Imaging Center, Hartwell Center, Center for Applied Bioinformatics, Center for Advanced Genomic Editing and the Neuroembryology core, and the Animal Resources Center at St. Jude Children’s Research Hospital (SJCRH) for help in image acquisition, sequencing, computational analysis, generating the *Insm1^HA^* allele, and animal care, respectively; Abbas Shirinifard for help in image quantification; Melvin DePamphilis and Jaime García-Añoveros for sharing mouse lines; David Ellison for providing medulloblastoma pathology images; James Morgan, Fabio Demontis, Stacey Ogden, David Solecki, and Michael Dyer for critical commenting of the manuscript. Microsoft Copilot and professional editing services from Life Science Editors were used for language editing. All scientific content was generated by the authors, who take full responsibility for the manuscript. This work was supported by National Institutes of Health grants R01NS119760 (to X.C.), P30CA021765 (to SJCRH Comprehensive Cancer Center core facilities), and American Lebanese Syrian Associated Charities.

## Author contributions

N.M.S. performed all histology associated experiments, image acquisition and quantification, and bulk RNA-sequencing; V.T. and J.N. performed single-cell RNA-seq experiment and data analysis; A.I. generated plasmid constructs and performed cell culture experiments; J.P. helped with mouse management and performed Western blots; Q.Z. and B.X. performed computational analysis of ATAC-seq and CUT&RUN data; X.C. conceptualized and supervised the project, performed co-immunoprecipitation, ATAC-seq, and CUT&RUN experiments, visualized the data, wrote the manuscript with input from other authors.

## Data, code, and materials availability

All high-throughput sequencing raw data have been deposited in the GEO (Gene Expression Omnibus) under accession codes GSE330837. This paper does not generate original code. Mouse models generated in this paper and any additional information required to reanalyze the data included in this paper are available from the corresponding author upon request.

## Supplementary Materials

**Fig. S1.**
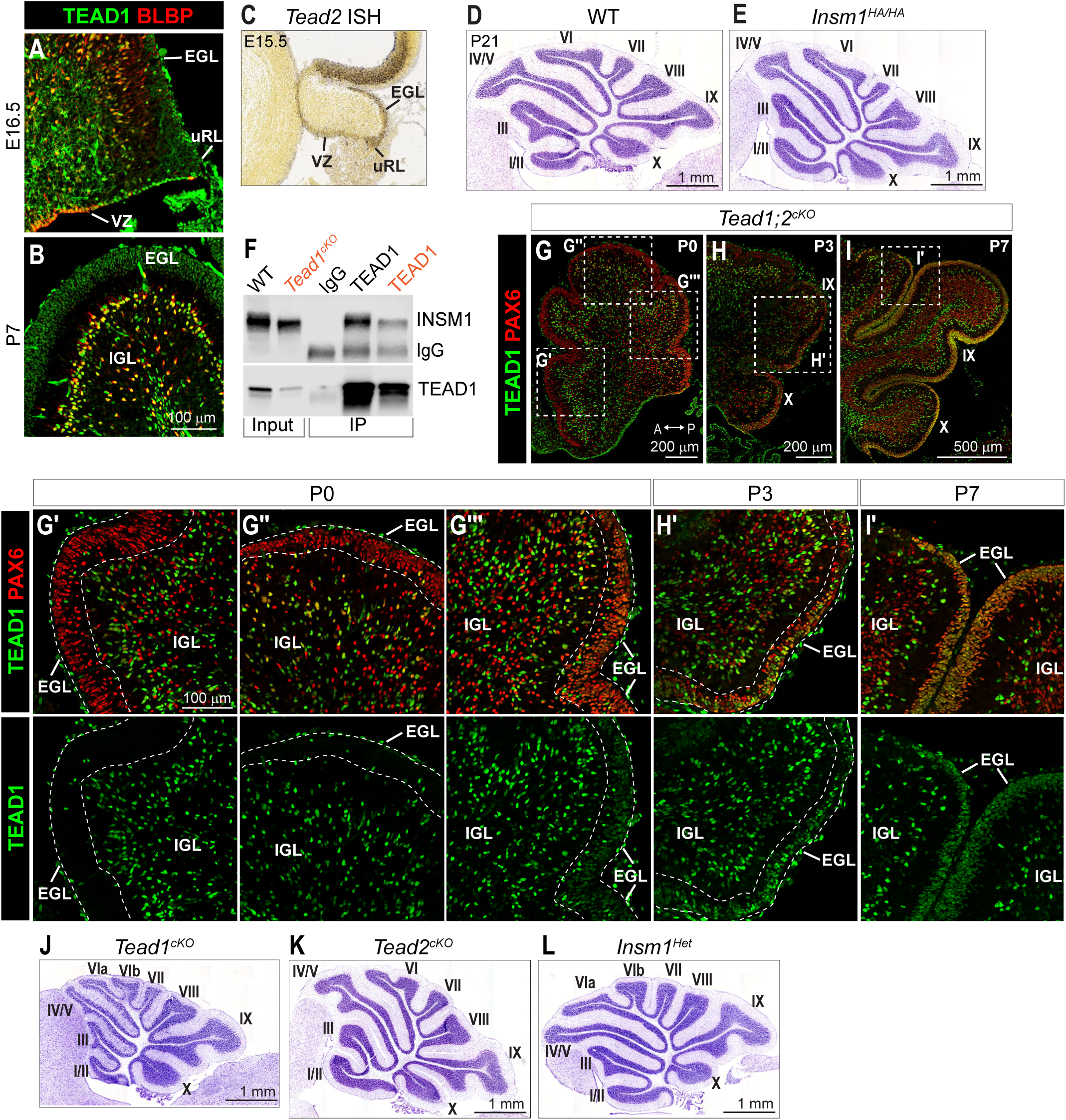
TEAD1/2 and INSM1 are essential for cerebellum development. (**A** and **B**) Co-immunostaining of wild-type (WT) cerebellum sections for TEAD1 and the glial marker BLBP. (**C**) In situ hybridization (ISH) detection of *Tead2* mRNA in E15.5 cerebellar primordium (Allen Brain Atlas). (**D** and **E**) Nissl staining of P21 cerebellum vermis sections. (**F**) Co-immunoprecipitation (co-IP) using IgG (negative control) or an anti-TEAD1 antibody from P7 WT or *Tead1^cKO^* (orange; negative control) cerebellar nuclear extracts. (**G** to **I′**) Immunostaining of P0, P3, and P7 *Tead1;2^cKO^* cerebella, showing loss of TEAD1 immunosignals in anterior and central lobules (G to G′′) but only sporadic loss in posteior lobules (IX and X; G′′′, H to I′). Boxed regions in G to I are shown at higher magnification in G′ to I′. (**J** to **L**) Nissl staining of P21 cerebella. VZ, ventricular zone; EGL, external granule layer; uRL, upper rhombic lip; IGL, internal granule layer.

**Fig. S2.**
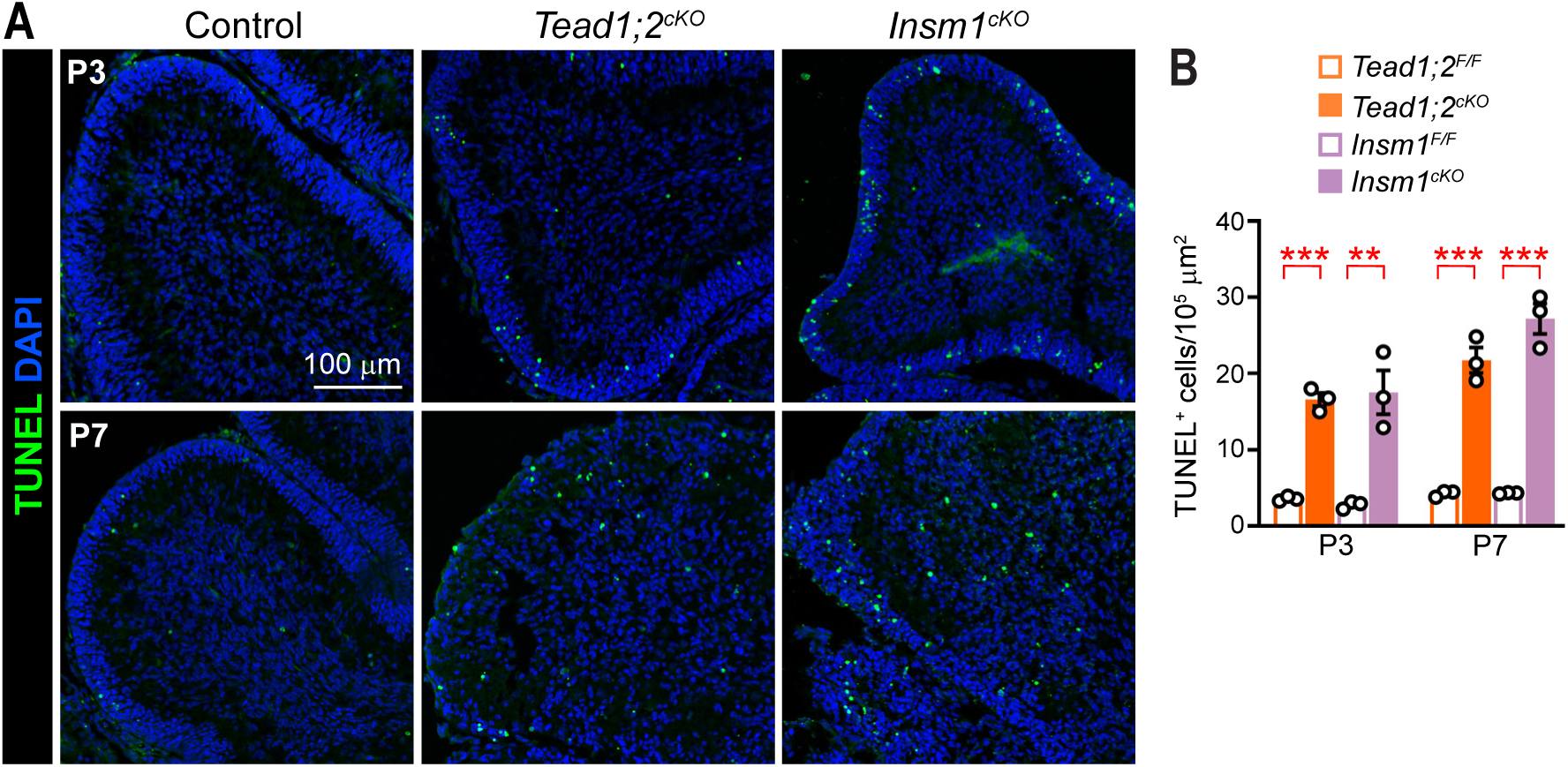
Loss of TEAD1/2 or INSM1 causes increased cell death during cerebellum development. (**A**) TUNEL labeling in P3 and P7 cerebellum sections. (**B**) Quantification of the number of TUNEL^+^ cells in lobules III–V. Each data point represents an individual animal. Values are mean ± SEM. Unpaired two-tailed *t*-test; **, *P* < 0.01; ***, *P* < 0.001.

**Fig. S3.**
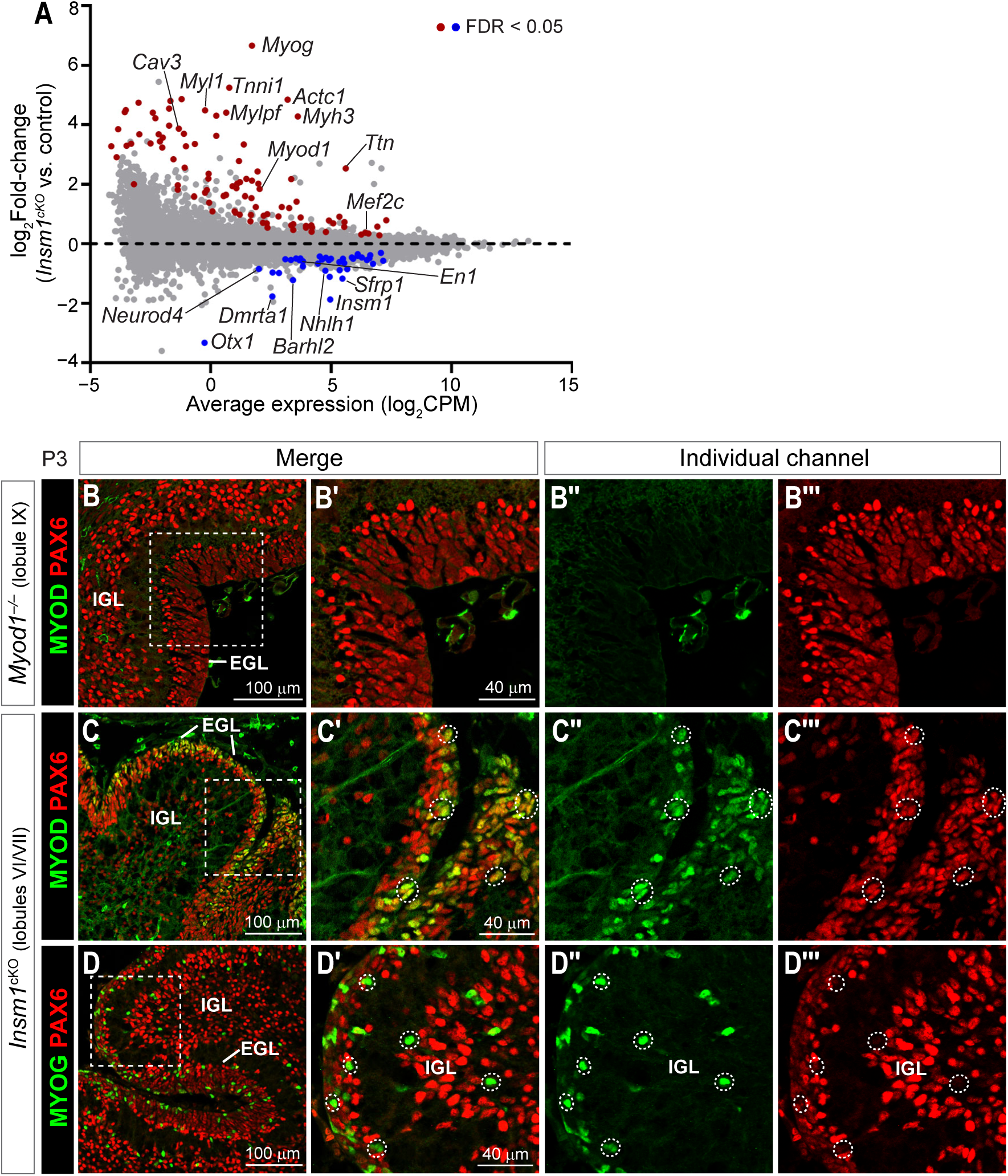
Loss of TEAD1/2 or INSM1 upregulates muscle genes. (**A**) MA plot showing gene expression changes in *Insm1^cKO^* cerebella. (**B** to **B′′′**) Co-immunostaining of P3 *Myod1^−/−^* cerebellum for PAX6 and MYOD. (**C** to **D′′′**) Co-immunostaining of P3 *Insm1^cKO^* cerebella for PAX6 and MYOD or MYOG. Boxed regions in B, C, and D are enlarged in B′, C′, and D′. Dotted circles highlight MYOD^+^ or MYOG^+^ cells. EGL, external granule layer; IGL, internal granule layer.

**Fig. S4.**
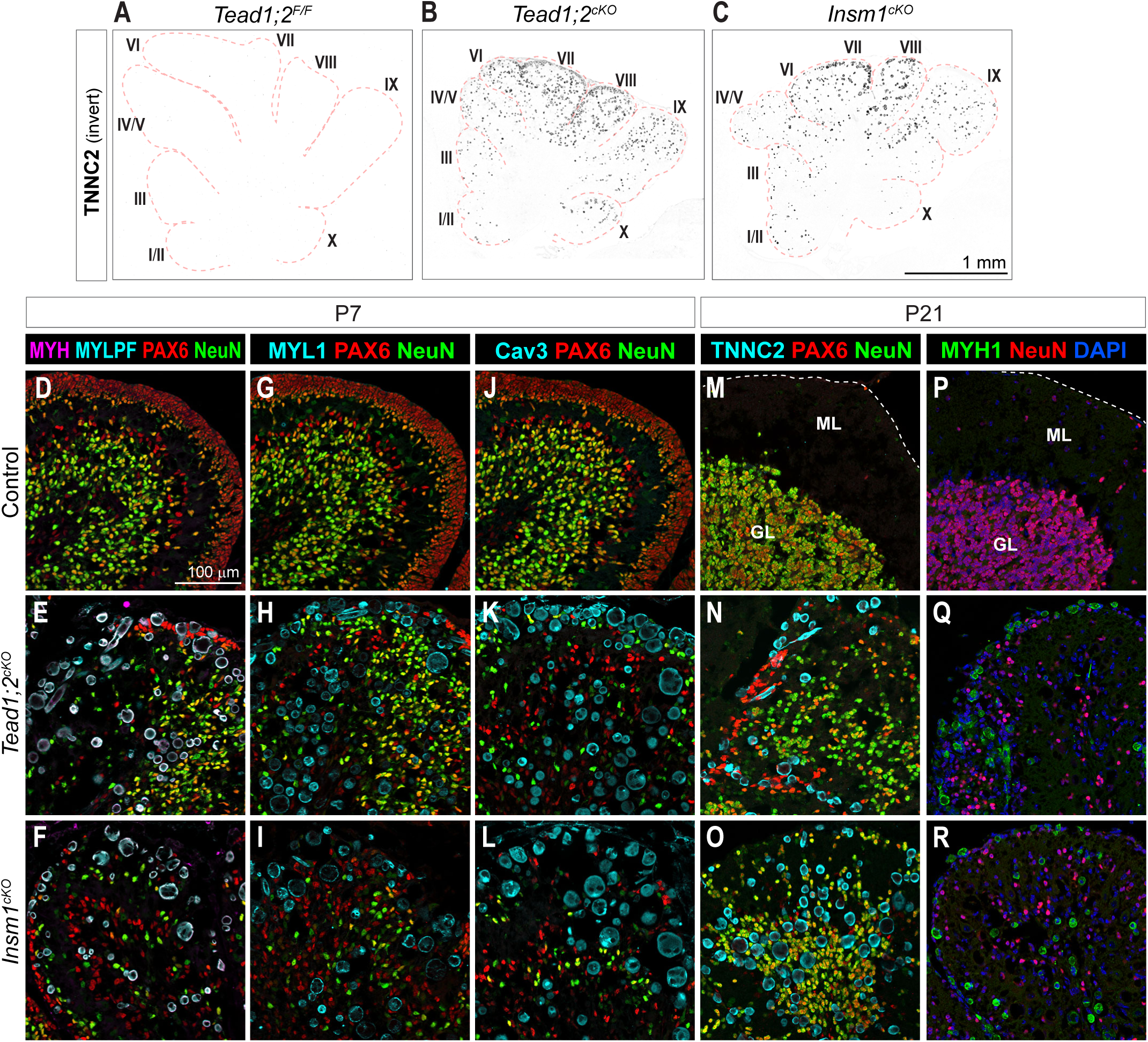
Expression of muscle structural proteins in *Tead1;2^cKO^* and *Insm1^cKO^* cerebella. (**A** to **C**) Immunostaining of P7 cerebella. Panels correspond to enlarged views of the TNNC2 (troponin C2) channel from Fig. 4, A, B, and E, respectively. Fluorescence signals were inverted for improved visualization. (**D** to **R**) Co-immunostaining of P7 and P21 cerebella for PAX6, NeuN, and muscle structural proteins myosin heavy chain (MYH), myosin light chain phosphorylatable (MYLPF), myosin light chain 1 (MYL1), Caveolin 3 (Cav3), and myosin heavy chain 1 (MYH1). The dashed lines in M and P demarcate tissue border. ML, molecular layer; GL, granule layer.

**Fig. S5.**
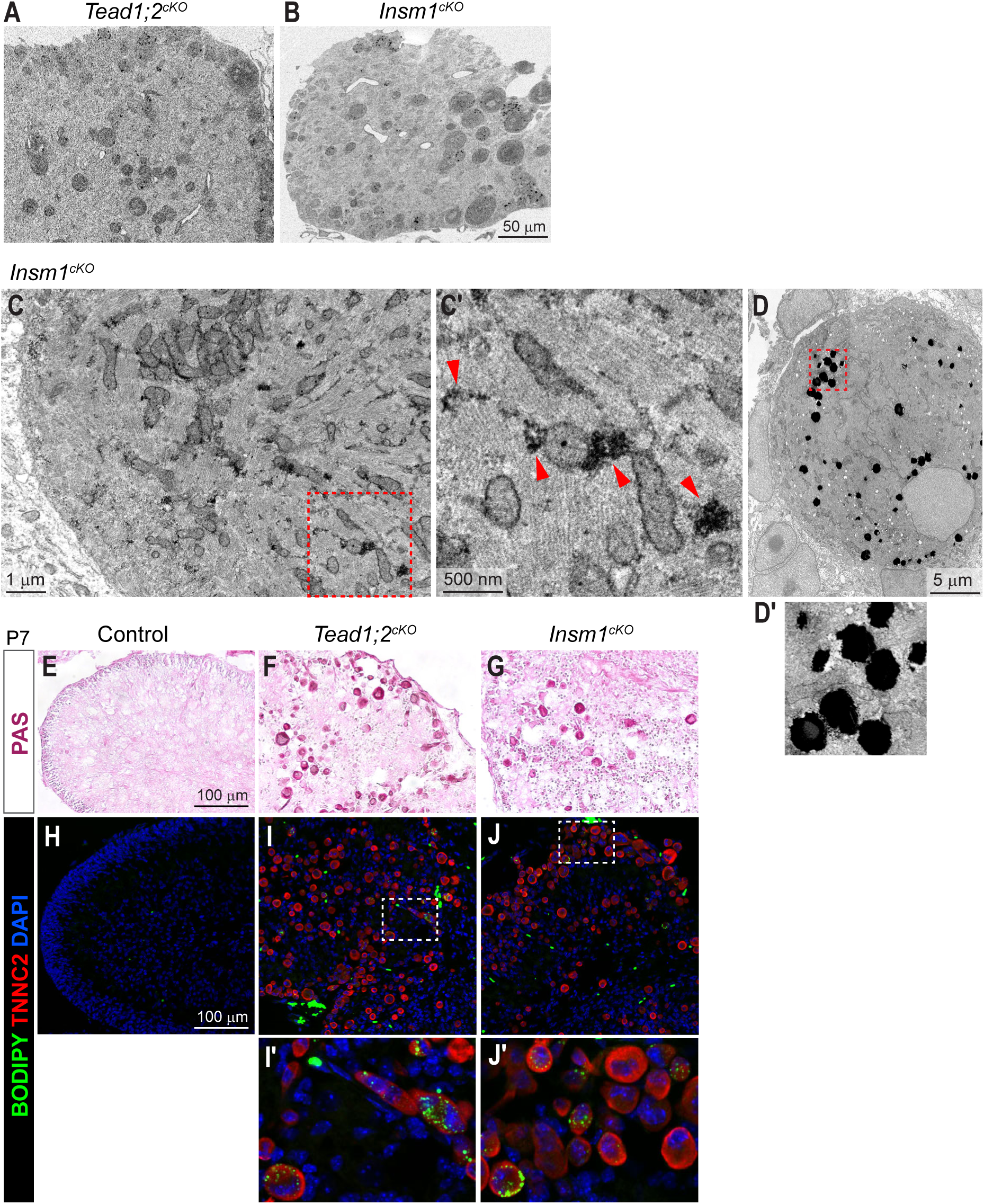
Characterization of muscle-like cells in *Tead1;2^cKO^* and *Insm1^cKO^* cerebella. (**A** and **B**) Low-magnification electron microscopy (EM) images of P7 mutant cerebellar sections. (**C** to **D′**) High-magnification EM images of P7 *Insm1^cKO^* cerebellar sections. Arrows, glycogen granules. (**E** to **G**) Periodic Acid-Schiff (PAS) staining of polysaccharides showing presence of glycogen (dark pink) in mutant cerebella. (**H** to **J′**) BODIPY staining combined with immunostaining showing lipid droplets in TNNC2^+^ cells in mutant cerebella. Boxed regions in C, D, I, and J are enlarged in C′, D′, I′, and J′, respectively.

**Fig. S6.**
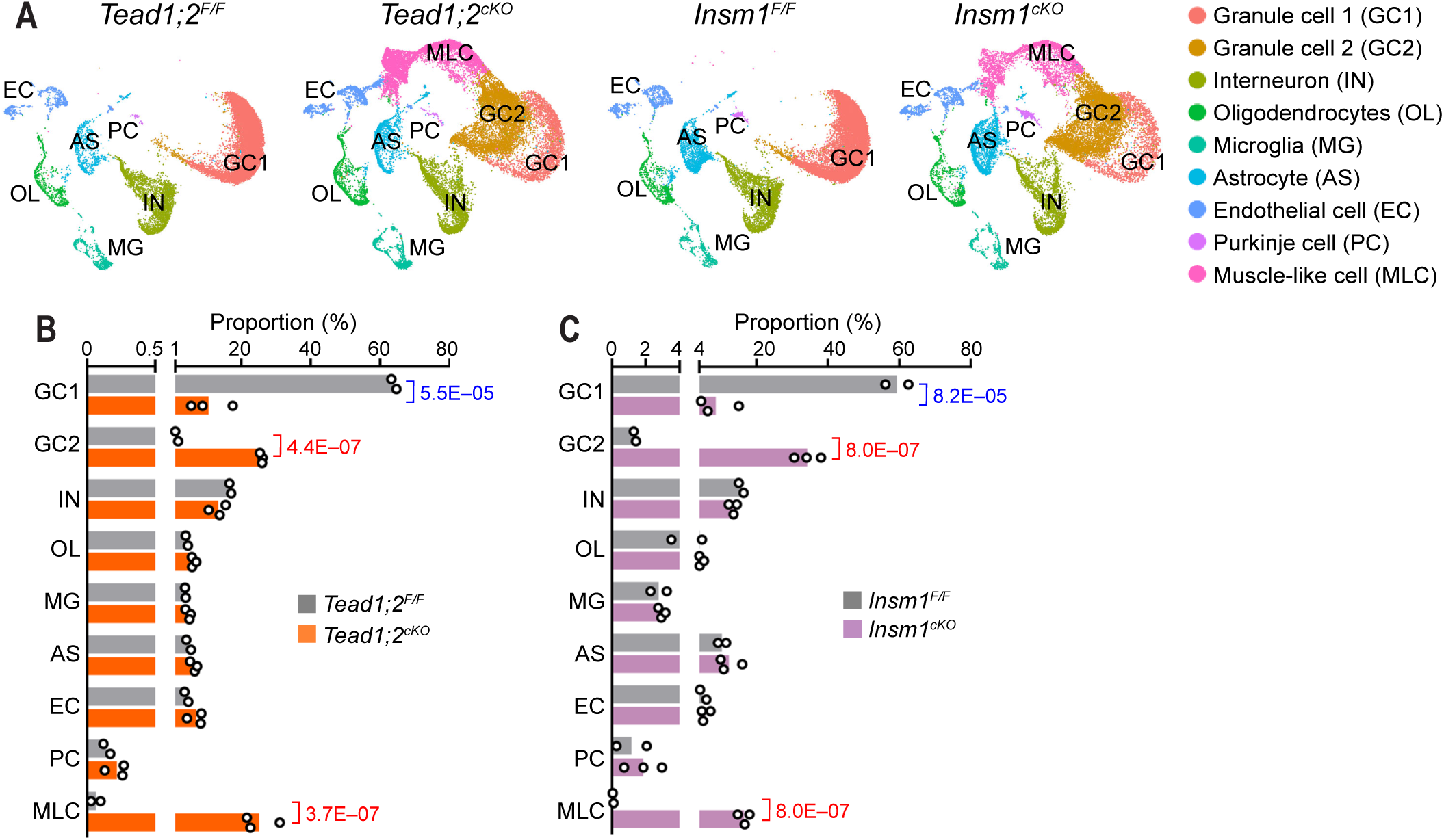
Single-cell RNA-sequencing analysis of P7 cerebella. (**A**) UMAP visualization and annotation of cerebellar cells based on genotype. *Tead1;2^F/F^* (*n* = 2 mice), 14,900 cells; *Tead1;2^cKO^* (*n* = 3), 22,783 cells; *Insm1^F/F^* (*n* = 2), 17,933 cells; *Insm1^cKO^* (*n* = 3), 18,173 cells. (**B** and **C**) Cell type proportions in each sample. Each data point represents an individual animal. Statistical test was performed using propeller. Values are false discovery rate (FDR). Unmarked comparions did not show a significant difference (FDR > 0.05).

**Fig. S7.**
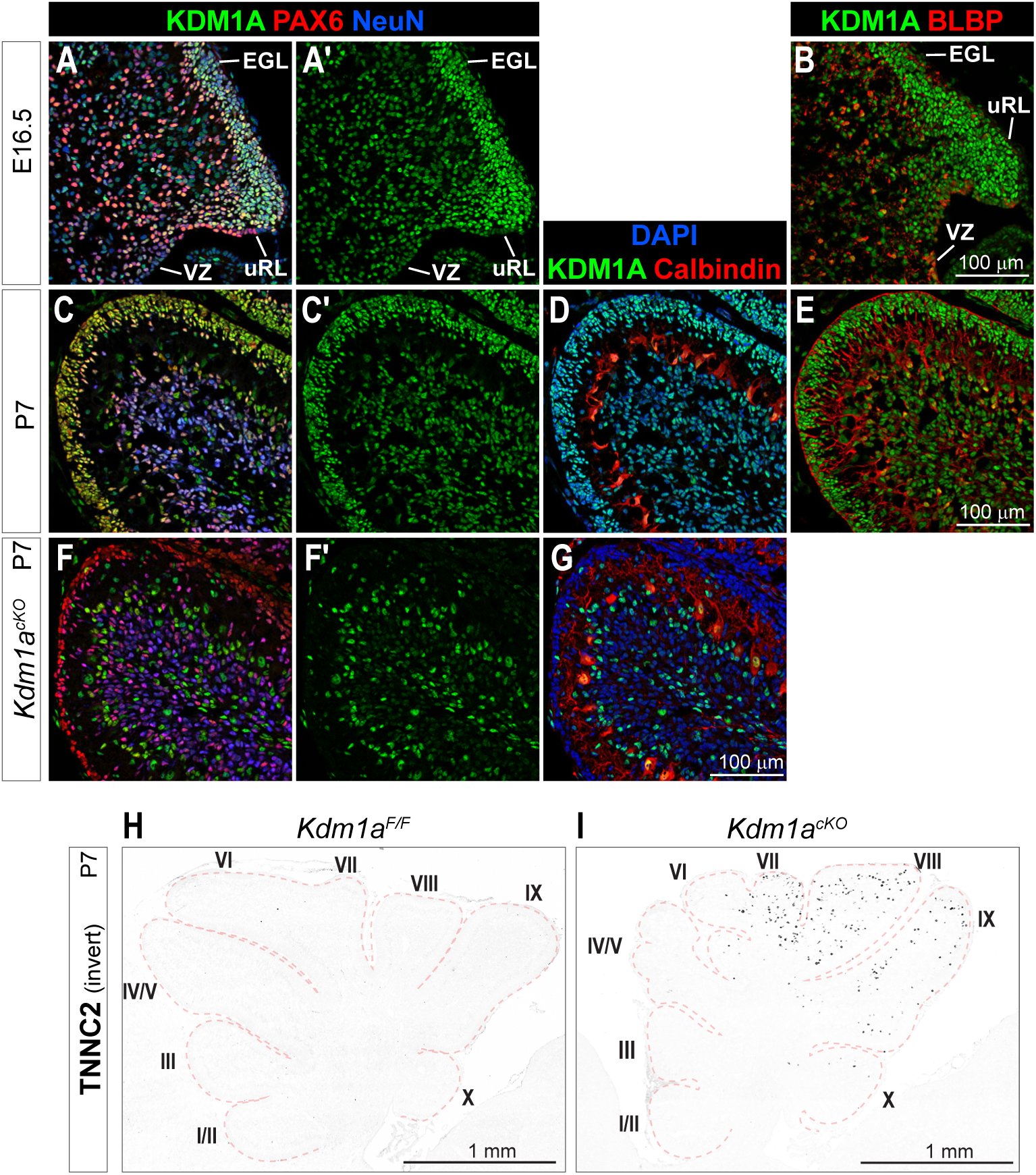
Characterization of KDM1A expression pattern and mutant phenotype during cerebellum development. (**A** to **E**) Co-immunostaining of E16.5 and P7 WT cerebella for KDM1A, the granule neuron lineage marker PAX6, the glial lineage marker BLBP, and the Purkinje cell marker Calbindin. (**F** to **G**) Co-immunostaining of P7 *Kdm1a^cKO^* cerebellum, showing loss of KDM1A immunosignal in PAX6^+^ cells. (**H** and **I**) The inverted and enlarged view of the TNNC2 channel from Fig. 5, F and G, respectively. VZ, ventricular zone; EGL, external granule layer; uRL, upper rhombic lip.

**Fig. S8.**
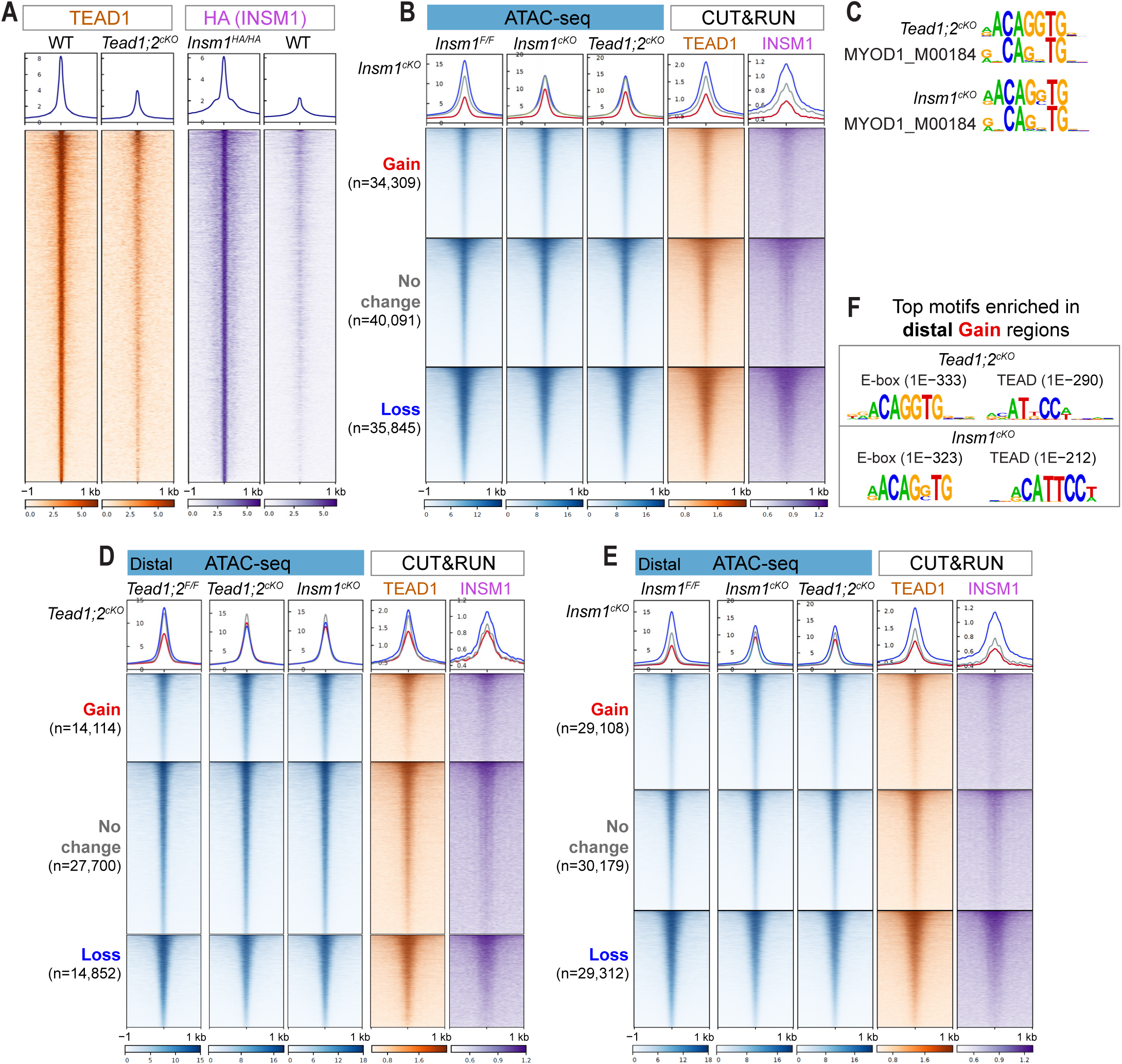
Changes in chromatin accessibility following the loss of TEAD1/2 or INSM1. (**A**) Profile graphs and heatmaps of TEAD1 and INSM1 genomic binding in P2 cerebella identified by CUT&RUN. (**B**) Profile graphs and heatmaps of chromatin accessibility (ATAC-seq) in P2 control and mutant cerebella, and TEAD1 and INSM1 genomic binding in P2 WT (*Insm1^HA/HA^* for INSM1) cerebella, grouped by their change in accessibility in *Insm1^cKO^* cerebella compared with littermate controls. Gain and Loss, FDR < 0.05, no fold-change (FC) cutoff; No change, |FC| < 1.05, FDR > 0.05. Profile graphs and heatmaps of ATAC-seq data were scaled based on the signals of no-change regions. One representative replicate of 2–4 biological replicates. (**C**) Alignment of the top enriched motif in the regions with gain of accessibility in mutant cerebella with MYOD1 consensus motif. (**D** and **E**) Profile graphs and heatmaps of chromatin accessibility of promoter-distal regions and TF binding grouped by their change in accessibility in *Tead1;2^cKO^* (D) and *Insm1^cKO^* (E) cerebella. (**F**) Top enriched motifs in distal regions with gain of accessibility in mutant cerebella compared with no-change regions, with *P* values.

**Fig. S9.**
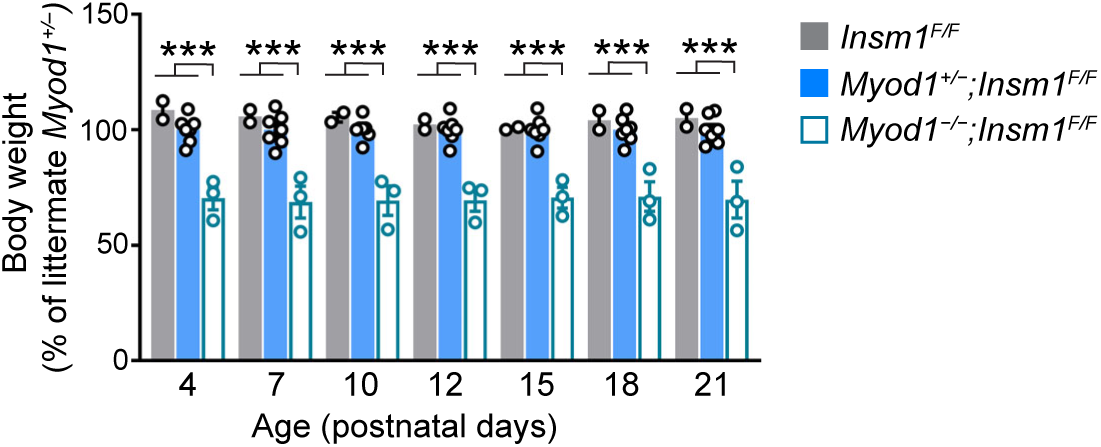
*Myod1* null mice show reduced body weight compared with littermate WT and heterozygous mice. Body weight of each mouse was normalized to the average weight of littermate heterozygous mice. Each data point represents an individual animal. Values are mean ± SEM. Unpaired two-tailed *t*-test; ***, *P* < 0.001.

Table S1. Bulk RNA-seq data.

Table S2. Single-cell RNA-seq signature genes.

Table S3. GREAT analysis of ATAC-seq differentially accessible regions: top 50 enriched gene sets.

## REFERENCES

1. M. Götz, M.-E. Torres-Padilla, Stem cells as role models for reprogramming and repair. Science 388, eadp2959 (2025).

2. T. Graf, T. Enver, Forcing cells to change lineages. Nature 462, 587–594 (2009).

3. J. M. Slack, Turning One Cell Type into Another. Current topics in developmental biology 117, 339–358 (2016).

4. H. Wang, Y. Yang, J. Liu, L. Qian, Direct cell reprogramming: approaches, mechanisms and progress. Nature Reviews Molecular Cell Biology 22, 410–424 (2021).

5. A. Davies, A. Zoubeidi, H. Beltran, L. A. Selth, The Transcriptional and Epigenetic Landscape of Cancer Cell Lineage Plasticity. Cancer Discovery 13, 1771–1788 (2023).

6. J. Padeken, S. P. Methot, S. M. Gasser, Establishment of H3K9-methylated heterochromatin and its functions in tissue differentiation and maintenance. Nature Reviews Molecular Cell Biology 23, 623–640 (2022).

7. R. L. McCarthy, J. Zhang, K. S. Zaret, Diverse heterochromatin states restricting cell identity and reprogramming. Trends in biochemical sciences 48, 513–526 (2023).

8. D. Nicetto, G. Donahue, T. Jain, T. Peng, S. Sidoli, L. Sheng, T. Montavon, J. S. Becker, J. M. Grindheim, K. Blahnik, B. A. Garcia, K. Tan, R. Bonasio, T. Jenuwein, K. S. Zaret, H3K9me3-heterochromatin loss at protein-coding genes enables developmental lineage specification. Science 363, 294–297 (2019).

9. S. P. Methot, J. Padeken, G. Brancati, P. Zeller, C. E. Delaney, D. Gaidatzis, H. Kohler, A. van Oudenaarden, H. Großhans, S. M. Gasser, H3K9me selectively blocks transcription factor activity and ensures differentiated tissue integrity. Nature Cell Biology 23, 1163–1175 (2021).

10. J. Brumbaugh, B. Di Stefano, K. Hochedlinger, Reprogramming: identifying the mechanisms that safeguard cell identity. Development 146, (2019).

11. Y. Zheng, D. Pan, The Hippo Signaling Pathway in Development and Disease. Dev Cell 50, 264–282 (2019).

12. N. Nishioka, K. Inoue, K. Adachi, H. Kiyonari, M. Ota, A. Ralston, N. Yabuta, S. Hirahara, R. O. Stephenson, N. Ogonuki, R. Makita, H. Kurihara, E. M. Morin-Kensicki, H. Nojima, J. Rossant, K. Nakao, H. Niwa, H. Sasaki, The Hippo signaling pathway components Lats and Yap pattern Tead4 activity to distinguish mouse trophectoderm from inner cell mass. Dev Cell 16, 398–410 (2009).

13. D. Jukam, B. Xie, J. Rister, D. Terrell, M. Charlton-Perkins, D. Pistillo, B. Gebelein, C. Desplan, T. Cook, Opposite feedbacks in the Hippo pathway for growth control and neural fate. Science 342, 1238016 (2013).

14. A. Sebé-Pedrós, Y. Zheng, I. Ruiz-Trillo, D. Pan, Premetazoan Origin of the Hippo Signaling Pathway. Cell Reports 1, 13–20 (2012).

15. J. E. Phillips, Y. Zheng, D. Pan, Assembling a Hippo: the evolutionary emergence of an animal developmental signaling pathway. Trends in biochemical sciences 49, 681–692 (2024).

16. J. H. A. Vissers, F. Froldi, J. Schröder, A. T. Papenfuss, L. Y. Cheng, K. F. Harvey, The Scalloped and Nerfin-1 Transcription Factors Cooperate to Maintain Neuronal Cell Fate. Cell Rep 25, 1561–1576.e1567 (2018).

17. P. Guo, C. H. Lee, H. Lei, Y. Zheng, K. D. Pulgar Prieto, D. Pan, Nerfin-1 represses transcriptional output of Hippo signaling in cell competition. eLife 8, (2019).

18. M. S. Lan, M. B. Breslin, Structure, expression, and biological function of INSM1 transcription factor in neuroendocrine differentiation. The FASEB Journal 23, 2024–2033 (2009).

19. J. Wu, A. Duggan, M. Chalfie, Inhibition of touch cell fate by egl-44 and egl-46 in C. elegans. Genes & Development 15, 789–802 (2001).

20. C. H. Perry, A. Lavado, V. Thulabandu, C. Ramirez, J. Paré, R. Dixit, A. Mishra, J. Yang, J. Yu, X. Cao, TEAD switches interacting partners along neural progenitor lineage progression to execute distinct functions. Genes Dev 39, 849–867 (2025).

21. X. Cao, S. L. Pfaff, F. H. Gage, YAP regulates neural progenitor cell number via the TEA domain transcription factor. Genes Dev 22, 3320–3334 (2008).

22. M. Grove, H. Kim, S. Pang, J. P. Amaya, G. Hu, J. Zhou, M. Lemay, Y.-J. Son, TEAD1 is crucial for developmental myelination, Remak bundles, and functional regeneration of peripheral nerves. eLife 13, e87394 (2024).

23. L. J. Hughes, R. Park, M. J. Lee, B. K. Terry, D. J. Lee, H. Kim, S. H. Cho, S. Kim, Yap/Taz are required for establishing the cerebellar radial glia scaffold and proper foliation. Dev Biol 457, 150–162 (2020).

24. G. G. Consalez, D. Goldowitz, F. Casoni, R. Hawkes, Origins, Development, and Compartmentation of the Granule Cells of the Cerebellum. Front Neural Circuits 14, 611841 (2020).

25. Allen Institute (2004). Allen Mouse Brain Atlas [dataset]. Available from mouse.brain-map.org. Allen Institute for Brain Science (2011).

26. G. Feng, P. Yi, Y. Yang, Y. Chai, D. Tian, Z. Zhu, J. Liu, F. Zhou, Z. Cheng, X. Wang, W. Li, G. Ou, Developmental stage-dependent transcriptional regulatory pathways control neuroblast lineage progression. Development 140, 3838–3847 (2013).

27. A. Duggan, T. Madathany, S. C. P. de Castro, D. Gerrelli, K. Guddati, J. García-Añoveros, Transient expression of the conserved zinc finger gene INSM1 in progenitors and nascent neurons throughout embryonic and adult neurogenesis. Journal of Comparative Neurology 507, 1497–1520 (2008).

28. M. S. Gierl, N. Karoulias, H. Wende, M. Strehle, C. Birchmeier, The Zinc-finger factor Insm1 (IA-1) is essential for the development of pancreatic β cells and intestinal endocrine cells. Genes & Development 20, 2465–2478 (2006).

29. A. B. Osipovich, Q. Long, E. Manduchi, R. Gangula, S. B. Hipkens, J. Schneider, T. Okubo, C. J. Stoeckert, Jr., S. Takada, M. A. Magnuson, Insm1 promotes endocrine cell differentiation by modulating the expression of a network of genes that includes Neurog3 and Ripply3. Development 141, 2939–2949 (2014).

30. V. Matei, S. Pauley, S. Kaing, D. Rowitch, K. W. Beisel, K. Morris, F. Feng, K. Jones, J. Lee, B. Fritzsch, Smaller inner ear sensory epithelia in Neurog 1 null mice are related to earlier hair cell cycle exit. Dev Dyn 234, 633–650 (2005).

31. N. Pan, I. Jahan, J. E. Lee, B. Fritzsch, Defects in the cerebella of conditional Neurod1 null mice correlate with effective Tg(Atoh1-cre) recombination and granule cell requirements for Neurod1 for differentiation. Cell Tissue Res 337, 407–428 (2009).

32. C. Atterton, A. Pelenyi, J. Jones, L. Currey, M. Al-Khalily, L. Wright, M. Doonan, D. Knight, N. D. Kurniawan, S. Walters, S. Thor, M. Piper, The Hippo effector TEAD1 regulates postnatal murine cerebellar development. Brain Struct Funct 230, 42 (2025).

33. P. S. Zammit, Function of the myogenic regulatory factors Myf5, MyoD, Myogenin and MRF4 in skeletal muscle, satellite cells and regenerative myogenesis. Semin Cell Dev Biol 72, 19–32 (2017).

34. J. Dey, A. M. Dubuc, K. D. Pedro, D. Thirstrup, B. Mecham, P. A. Northcott, X. Wu, D. Shih, S. J. Tapscott, M. LeBlanc, M. D. Taylor, J. M. Olson, MyoD Is a Tumor Suppressor Gene in Medulloblastoma. Cancer Research 73, 6828–6837 (2013).

35. S. Schiaffino, A. C. Rossi, V. Smerdu, L. A. Leinwand, C. Reggiani, Developmental myosins: expression patterns and functional significance. Skelet Muscle 5, 22 (2015).

36. K. Gupta, S. Jogunoori, A. Satapathy, P. Salunke, N. Kumar, B. D. Radotra, R. K. Vasishta, Medulloblastoma with myogenic and/or melanotic differentiation does not align immunohistochemically with the genetically defined molecular subgroups. Hum Pathol 75, 26–33 (2018).

37. M. Sepp, K. Leiss, F. Murat, K. Okonechnikov, P. Joshi, E. Leushkin, L. Spänig, N. Mbengue, C. Schneider, J. Schmidt, N. Trost, M. Schauer, P. Khaitovich, S. Lisgo, M. Palkovits, P. Giere, L. M. Kutscher, S. Anders, M. Cardoso-Moreira, I. Sarropoulos, S. M. Pfister, H. Kaessmann, Cellular development and evolution of the mammalian cerebellum. Nature 625, 788–796 (2024).

38. J. E. Welcker, L. R. Hernandez-Miranda, F. E. Paul, S. Jia, A. Ivanov, M. Selbach, C. Birchmeier, Insm1 controls development of pituitary endocrine cells and requires a SNAG domain for function and for recruitment of histone-modifying factors. Development 140, 4947–4958 (2013).

39. X. de Martin, R. Sodaei, G. Santpere, Mechanisms of Binding Specificity among bHLH Transcription Factors. International Journal of Molecular Sciences 22, 9150 (2021).

40. R. L. Davis, H. Weintraub, A. B. Lassar, Expression of a single transfected cDNA converts fibroblasts to myoblasts. Cell 51, 987–1000 (1987).

41. H. Weintraub, S. J. Tapscott, R. L. Davis, M. J. Thayer, M. A. Adam, A. B. Lassar, A. D. Miller, Activation of muscle-specific genes in pigment, nerve, fat, liver, and fibroblast cell lines by forced expression of MyoD. Proc Natl Acad Sci U S A 86, 5434–5438 (1989).

42. J. Choi, M. L. Costa, C. S. Mermelstein, C. Chagas, S. Holtzer, H. Holtzer, MyoD converts primary dermal fibroblasts, chondroblasts, smooth muscle, and retinal pigmented epithelial cells into striated mononucleated myoblasts and multinucleated myotubes. Proc Natl Acad Sci U S A 87, 7988–7992 (1990).

43. F. C. Wardle, Master control: transcriptional regulation of mammalian Myod. J Muscle Res Cell Motil 40, 211–226 (2019).

44. M. A. Rudnicki, P. N. Schnegelsberg, R. H. Stead, T. Braun, H. H. Arnold, R. Jaenisch, MyoD or Myf-5 is required for the formation of skeletal muscle. Cell 75, 1351–1359 (1993).

45. V. Thulabandu, X. Cao, Insulinoma-associated 1 promotes neurogenic proliferation of cortical basal progenitors but is largely dispensable for projection neuron production. bioRxiv. 2026 (10.64898/2026.04.02.716139).

46. L. M. Farkas, C. Haffner, T. Giger, P. Khaitovich, K. Nowick, C. Birchmeier, S. Paabo, W. B. Huttner, Insulinoma-associated 1 has a panneurogenic role and promotes the generation and expansion of basal progenitors in the developing mouse neocortex. Neuron 60, 40–55 (2008).

47. T. Wiwatpanit, S. M. Lorenzen, J. A. Cantú, C. Z. Foo, A. K. Hogan, F. Márquez, J. C. Clancy, M. J. Schipma, M. A. Cheatham, A. Duggan, J. García-Añoveros, Trans-differentiation of outer hair cells into inner hair cells in the absence of INSM1. Nature 563, 691–695 (2018).

48. M. A. Forbes-Osborne, S. G. Wilson, A. C. Morris, Insulinoma-associated 1a (Insm1a) is required for photoreceptor differentiation in the zebrafish retina. Dev Biol 380, 157–171 (2013).

49. A. H. Cleveland, D. Malawsky, M. Churiwal, C. Rodriguez, F. Reed, M. Schniederjan, J. E. Velazquez Vega, I. Davis, T. R. Gershon, PRC2 disruption in cerebellar progenitors produces cerebellar hypoplasia and aberrant myoid differentiation without blocking medulloblastoma growth. Acta Neuropathol Commun 11, 8 (2023).

50. S. Joshi, G. Davidson, S. Le Gras, S. Watanabe, T. Braun, G. Mengus, I. Davidson, TEAD transcription factors are required for normal primary myoblast differentiation in vitro and muscle regeneration in vivo. PLoS Genet 13, e1006600 (2017).

51. T. Homma, A. Hemmi, T. Ohta, Y. Kusumi, A. Yoshino, H. Hao, A rare case of a pineoblastoma with a rhabdomyoblastic component. Neuropathology 37, 227–232 (2017).

52. Y. Wei, Q. Qin, C. Yan, M. N. Hayes, S. P. Garcia, H. Xi, D. Do, A. H. Jin, T. C. Eng, K. M. McCarthy, A. Adhikari, M. L. Onozato, D. Spentzos, G. P. Neilsen, A. J. Iafrate, L. H. Wexler, A. D. Pyle, M. L. Suvà, F. Dela Cruz, L. Pinello, D. M. Langenau, Single-cell analysis and functional characterization uncover the stem cell hierarchies and developmental origins of rhabdomyosarcoma. Nature Cancer 3, 961–975 (2022).

53. M. Hayano, J. H. Sung, A. R. Mastri, E. G. Hill, Striated muscle in the pineal gland of swine. J Neuropathol Exp Neurol 35, 613–621 (1976).

54. B. J. Diehl, Occurrence and regional distribution of striated muscle fibers in the rat pineal gland. Cell Tissue Res 190, 349–355 (1978).

55. D. Arendt, The evolution of cell types in animals: emerging principles from molecular studies. Nat Rev Genet 9, 868–882 (2008).

56. D. Arendt, E. Benito-Gutierrez, T. Brunet, H. Marlow, Gastric pouches and the mucociliary sole: setting the stage for nervous system evolution. Philos Trans R Soc Lond B Biol Sci 370, (2015).

57. K. Seipel, N. Yanze, V. Schmid, Developmental and evolutionary aspects of the basic helix-loop-helix transcription factors Atonal-like 1 and Achaete-scute homolog 2 in the jellyfish. Dev Biol 269, 331–345 (2004).

58. M. A. Kerenyi, Z. Shao, Y. J. Hsu, G. Guo, S. Luc, K. O’Brien, Y. Fujiwara, C. Peng, M. Nguyen, S. H. Orkin, Histone demethylase Lsd1 represses hematopoietic stem and progenitor cell signatures during blood cell maturation. eLife 2, e00633 (2013).

59. M. A. Rudnicki, T. Braun, S. Hinuma, R. Jaenisch, Inactivation of MyoD in mice leads to up-regulation of the myogenic HLH gene Myf-5 and results in apparently normal muscle development. Cell 71, 383–390 (1992).

60. K. J. Kaneko, M. J. Kohn, C. Liu, M. L. DePamphilis, Transcription factor TEAD2 is involved in neural tube closure. Genesis 45, 577–587 (2007).

61. C. Genoud, B. Titze, A. Graff-Meyer, R. W. Friedrich, Fast Homogeneous En Bloc Staining of Large Tissue Samples for Volume Electron Microscopy. Front Neuroanat 12, 76 (2018).

62. J. Walton, Lead aspartate, an en bloc contrast stain particularly useful for ultrastructural enzymology. J Histochem Cytochem 27, 1337–1342 (1979).

63. F. Krueger, F. James, P. Ewels, E. Afyounian, B. Schuster-Boeckler. (Zenodo, 2021).

64. A. Dobin, T. R. Gingeras, Mapping RNA-seq reads with STAR. Current protocols in bioinformatics 51, 11.14. 11–11.14. 19 (2015).

65. B. Li, C. N. Dewey, RSEM: accurate transcript quantification from RNA-Seq data with or without a reference genome. BMC bioinformatics 12, 323 (2011).

66. C. W. Law, Y. Chen, W. Shi, G. K. Smyth, voom: Precision weights unlock linear model analysis tools for RNA-seq read counts. Genome Biol 15, R29 (2014).

67. B. Phipson, C. B. Sim, E. R. Porrello, A. W. Hewitt, J. Powell, A. Oshlack, propeller: testing for differences in cell type proportions in single cell data. Bioinformatics 38, 4720–4726 (2022).

68. H. Li, R. Durbin, Fast and accurate short read alignment with Burrows-Wheeler transform. Bioinformatics 25, 1754–1760 (2009).

69. H. Li, B. Handsaker, A. Wysoker, T. Fennell, J. Ruan, N. Homer, G. Marth, G. Abecasis, R. Durbin, The Sequence Alignment/Map format and SAMtools. Bioinformatics 25, 2078–2079 (2009).

70. A. R. Quinlan, I. M. Hall, BEDTools: a flexible suite of utilities for comparing genomic features. Bioinformatics 26, 841–842 (2010).

71. R. M. Kuhn, D. Haussler, W. J. Kent, The UCSC genome browser and associated tools. Brief Bioinform 14, 144–161 (2013).

72. J. T. Robinson, H. Thorvaldsdóttir, W. Winckler, M. Guttman, E. S. Lander, G. Getz, J. P. Mesirov, Integrative genomics viewer. Nat Biotechnol 29, 24–26 (2011).

73. Y. Zhang, T. Liu, C. A. Meyer, J. Eeckhoute, D. S. Johnson, B. E. Bernstein, C. Nusbaum, R. M. Myers, M. Brown, W. Li, X. S. Liu, Model-based analysis of ChIP-Seq (MACS). Genome Biol 9, R137 (2008).

74. R. K. Dale, B. S. Pedersen, A. R. Quinlan, Pybedtools: a flexible Python library for manipulating genomic datasets and annotations. Bioinformatics 27, 3423–3424 (2011).

75. Y. Zhang, G. Parmigiani, W. E. Johnson, ComBat-seq: batch effect adjustment for RNA-seq count data. NAR Genom Bioinform 2, lqaa078 (2020).

76. M. D. Robinson, D. J. McCarthy, G. K. Smyth, edgeR: a Bioconductor package for differential expression analysis of digital gene expression data. Bioinformatics 26, 139–140 (2010).

77. M. E. Ritchie, B. Phipson, D. Wu, Y. Hu, C. W. Law, W. Shi, G. K. Smyth, limma powers differential expression analyses for RNA-sequencing and microarray studies. Nucleic Acids Res 43, e47 (2015).

78. A. Frankish, M. Diekhans, A. M. Ferreira, R. Johnson, I. Jungreis, J. Loveland, J. M. Mudge, C. Sisu, J. Wright, J. Armstrong, I. Barnes, A. Berry, A. Bignell, S. Carbonell Sala, J. Chrast, F. Cunningham, T. Di Domenico, S. Donaldson, I. T. Fiddes, C. García Girón, J. M. Gonzalez, T. Grego, M. Hardy, T. Hourlier, T. Hunt, O. G. Izuogu, J. Lagarde, F. J. Martin, L. Martínez, S. Mohanan, P. Muir, F. C. P. Navarro, A. Parker, B. Pei, F. Pozo, M. Ruffier, B. M. Schmitt, E. Stapleton, M. M. Suner, I. Sycheva, B. Uszczynska-Ratajczak, J. Xu, A. Yates, D. Zerbino, Y. Zhang, B. Aken, J. S. Choudhary, M. Gerstein, R. Guigó, T. J. P. Hubbard, M. Kellis, B. Paten, A. Reymond, M. L. Tress, P. Flicek, GENCODE reference annotation for the human and mouse genomes. Nucleic Acids Res 47, D766–d773 (2019).

79. C. Y. McLean, D. Bristor, M. Hiller, S. L. Clarke, B. T. Schaar, C. B. Lowe, A. M. Wenger, G. Bejerano, GREAT improves functional interpretation of cis-regulatory regions. Nat Biotechnol 28, 495–501 (2010).

80. Y. Tanigawa, E. S. Dyer, G. Bejerano, WhichTF is functionally important in your open chromatin data? PLoS Comput Biol 18, e1010378 (2022).

81. S. Heinz, C. Benner, N. Spann, E. Bertolino, Y. C. Lin, P. Laslo, J. X. Cheng, C. Murre, H. Singh, C. K. Glass, Simple combinations of lineage-determining transcription factors prime cis-regulatory elements required for macrophage and B cell identities. Mol Cell 38, 576–589 (2010).

82. M. P. Meers, T. D. Bryson, J. G. Henikoff, S. Henikoff, Improved CUT&RUN chromatin profiling tools. eLife 8, (2019).

83. S. Xu, S. Grullon, K. Ge, W. Peng, Spatial clustering for identification of ChIP-enriched regions (SICER) to map regions of histone methylation patterns in embryonic stem cells. Methods in molecular biology (Clifton, N.J.) 1150, 97–111 (2014).

